# Adaptive variation in drought-related traits across southern and central European white oak (*Quercus* sect. *Quercus*) populations

**DOI:** 10.64898/2026.08.23.746538

**Authors:** DM Leigh, P Acar, C Blyth, SA Jansen, A Kremer, A Piotti, V Popovic, R Graf, NS McNamara, V Vitali, M Saurer, O Mavi İdman, Z Kaya, C Neophytou, C Rellstab

## Abstract

European white oaks grow from the Mediterranean coast to Southern Scandinavia, a huge environmental gradient that has likely fostered environmental adaptation. In the face of climate change, leveraging adaptations through assisted gene flow could help improve drought tolerance and maintain forest health, but requires an understanding of the species-specific patterns of adaptation to be successful at the target location. In this study, three common gardens were established in Switzerland, Türkiye, and Austria for two European white oak species (*Quercus robur*, and *Q. pubescens)* using provenances from Central and Southern Europe. Almost 900 oak seedlings were measured at key water-use efficiency and life history traits for their first two year of life and genotyped with low coverage whole-genome sequencing. Trait heritability and environmental adaptation were then explored through pedigree-free animal models, while the genomic architecture of traits was mapped using a genome wide association study (“GWAS”). Across the species, the heritability of measured traits was moderate to high, but common garden had a strong impact, signalling an environmental effect on the phenotype. Adaptation to precipitation seasonality was detected in key productivity and growth traits for both species, but had a small effect on absolute trait values. The GWAS identified a striking 150 kbp association in the Cyclic Nucleotide-Gated Ion Channel gene family with leaf δ^13^C values. This gene family is involved in stomata opening and likely impacts the intrinsic water use efficiency under stress. Together, the strong signals of phenotypic plasticity and rather weak signals of climatic adaptation in seedlings suggest that assisted gene flow in these two white oaks is relevant only for highly drought-sensitive populations, if conducted managers should focus on seeds sources with high precipitation seasonality and smaller leaf sizes.

## Introduction

Human-mediated climate change is leading to an increase in drought frequency and intensity across the globe (Chiang et al., 2021; Pokhrel et al., 2021). This repatterning of water availability is already impacting European forests through increased canopy damage and tree mortality (Allen et al., 2010, 2015; Senf et al., 2020). The long-term ecological and economic consequences of drought-related damage are forecast to be severe, especially because trees are keystone species which underpin high value ecosystem services and large ecological networks (Hanewinkel et al., 2013). To preserve these roles, it is vital that mitigation strategies are put in place to help reduce drought damage and maintain forest health (Aitken & Bemmels, 2016; Aitken & Whitlock, 2013).

Many common European tree species have vast geographical distributions, which naturally encompass large environmental gradients that include different degrees of historical exposure to drought. In regions where drought has been historically common, adaptations to drought are likely to have arisen (e.g. in *Quercus suber,* Morillas et al. 2024). It has been proposed that forest managers could minimize climate change-mediated damage by transferring and using reproductive material from source populations that are locally adapted to drought to regions newly encountering drought (termed ‘assisted gene flow’; Aitken & Bemmels, 2016; Aitken & Whitlock, 2013). Once established, the transferred genotypes will be better able to cope with drought relative to the resident genotypes. Over time, trees will interbreed and thus spread drought-adapted alleles leading to a general relative increase in forest health (Aitken & Bemmels, 2016; Aitken & Whitlock, 2013). Unlike strategies that move species out of their current distribution (termed ‘assisted migration’), assisted gene flow maintains the local ecological communities dependent on specific tree species, reducing the harmful ecological repercussions of climate-change mediated tree damage.

It is often colloquially assumed that seeds or trees selected for assisted gene flow should be moved from southern latitudes that have higher mean temperatures and lower annual rainfall (i.e., historical exposure to drought), to northern latitudes where these conditions are increasing under climate change. However, while several studies on tree species have identified broad continental scale differences of adaptations (Isaac-Renton et al., 2018; Rabarijaona et al., 2022), many trees species may also be highly plastic and innately drought-tolerant throughout their range (Candido-Ribeiro & Aitken, 2024). For such species, assisted gene flow may be unnecessary, have limited returns, or carry too high associated risks. These risks include phenological mismatches due to the natural seasonal differences across latitudes (Le Provost et al., 2023), photoperiod differences (Aitken & Bemmels, 2016), transfer of pathogens within seeds (Franić et al., 2019), or reducing productivity because of missing co-adaptation to local symbionts or resistance (Chen et al., 2020). There is also the risk that transfer may be counterproductive if climatic adaptation is highly regional with specific gene networks controlling adaptations in different areas (Browne et al., 2019; Mead et al., 2019; West et al., 2025). Collectively, it is important to first understand if environmental adaptation is present and describe its patterns in a species before implementing assisted gene flow programs.

European white oaks (*Quercus* sect. *Quercus*) are keystone species that support species-rich communities and underpin high-value ecosystem services (Mitchell et al., 2019). Due to the ecological, economic and cultural value of white oaks (Mitchell et al., 2019), as well as their high drought tolerance (Schotte et al., 2026; Wang et al., 2025), oak focused forestry is of high importance throughout Europe. White oaks have broad geographical ranges (EUFORGEN, 2025), which encompass an array of environmental conditions, including a particularly wide range of soil moisture conditions relative to other hardwood forest trees (Eaton et al., 2016; Rellstab, Zoller, et al., 2016). Environmental adaptation facilitates the large environmental niche of oak species and several studies have begun to identify or map adaptation through common gardens (e.g. bud burst, chlorophyll traits, Bantis et al. 2020; height, Mátyás 2021; height and survival, Bogdan, et al., 2017; growth traits, Martínez-Sancho et al. 2025; specific leaf area “SLA”, Morcillo et al. 2020; aeedling growth, Arend et al. 2011), genome-wide association analysis “GWAS” (e.g. wood related traits and growth traits, Lobo et al., 2025; Pålsson et al. 2026) and genotype environment association “GEA” analyses (Rellstab, Zoller, et al., 2016; Temunović et al., 2020).

The European white oaks *Q. robur* L. (pedunculate/English oak)*, Q. pubescens* Willd. (pubescent/downy oak) and *Q. petraea* (Matt.) Liebl. (sessile oak) are part of a widespread thermophilic species complex, showing weak species-level differentiation (Denk & Grimm, 2010) and secondary contact (Leroy et al., 2020). Typically, they are considered drought-resilient species because of their anisohydric leaves that allow for internal water balance adjustment to postpone stomatal closure under water stress, as well as their deep roots that allow them to access deep water reserves (Schotte et al., 2026; Wang et al., 2025). These characteristics have thus far buffered the species in Northern Europe from the extreme-drought damage seen in other dominant trees (Schotte et al., 2026; Wang et al., 2025). Drought-related adaptations have been identified in *Q. robur* in intrinsic water-use efficiency (Niemczyk et al., 2025) and through GEA associations with precipitation (Rellstab, Zoller, et al., 2016). However, drought-mediated oak growth decline and mortality has begun to increase in *Q. robur* (Gosling et al., 2024; Neumair et al., 2022; Truffaut et al., 2017), particularly in southern Europe in conjunction with comorbidities from fungal disease or insect pests (e.g. Colangelo et al. 2018; Gosling et al. 2024). Damage severity often varies within populations suggesting partial genetic control of drought response and resilience (e.g. *Q. robur* Colangelo et al. 2018; Niemczyk et al. 2025). Importantly, drought mortality trends are also not equal across the species. Mortality trends are more variable in *Q. pubescens*, which has a more southern distribution and drought-associated niche (Neumair et al., 2022; Rellstab, Zoller et al., 2016). It is generally considered to be a suitable species for climate smart forestry (Wöhlbrandt et al., 2026), however it is less exploited by foresters and substantially less studied than *Q. robur*, thus the patterns of adaptation in the species remain unclear. A comprehensive understanding of drought-related adaptation is thus needed for both species, particularly for populations in South and Central Europe where the projected increases in drought are extreme (Hari et al., 2020; Sonny et al., 2026) and because of the putative value of southern provenances as sources for assisted gene flow.

Combining common garden experiments with genomic analyses represents a powerful avenue for gaining insight into oak drought adaptation (de Villemereuil et al., 2016), because it can be used to estimate trait heritability, identify the presence of climatic adaptation, as well as genomic architecture of relevant traits. Combining common gardens with low coverage whole-genome sequencing (lcWGS) is further promising, because it is cost effective, allowing for the large samples sizes and individual identification (Korneliussen et al., 2014). Because each individual is sequenced to a reduced depth (typically <5x coverage), genotype likelihoods (GLs) are used and each individual receives a likelihood of each three possible genotypes (e.g. AA/Aa/aa) at a polymorphic locus. GLs incorporate coverage information, allele frequencies errors, and uncertainty (Korneliussen et al., 2014). Specialized analytical tools that can process these GL are required (Korneliussen et al., 2014; Skotte et al., 2013). To our knowledge, a lcWGS approach has yet to be extensively leveraged for heritability estimations but offers promising power and resolution.

In this study, we coupled common gardens focused on southern and central European populations of *Q. robur* and *Q. pubescens* with lcWGS to characterise climate adaptation in key seedling life-history and drought-related traits. Specifically, we used three common gardens of seedlings grown for two years to estimate seedling trait narrow-sense heritability and maternal effects with a pedigree-free animal model. After controlling for baseline relatedness, we searched for climate adaptation by looking for associations between trait values and climate of origin (focusing on precipitation, temperature, and precipitation variability). We then built on this analysis by using a GWAS to map trait architecture to help further studies understand the genes of importance in drought adaptation.

## Materials and Methods

### Seed collection and establishment of common gardens

In autumn of 2020 and 2021, acorns were collected for three white oak species (*Q. pubescens; Q. robur; Q. petraea*) from Central and Southern Europe. Acorn collection sites were chosen from three different geographical scales: local, regional, and continental. Sites at the local scale consisted of a pair of oak stands <10 km apart with contrasting soil moisture conditions (persistently wet vs dry) to capture local differences in edaphic drought stress. Sites at the regional scale consisted of replicated pairs. Continental sites were latitudinal extremes, which were not replicated due to inherent range constraints, and were chosen to encompass large differences in drought history (Figure 1). We focused on natural, non-mixed stands for seed provenances. This sampling was achieved in *Q. pubescens* (Figure 1). However, this was not possible in *Q. robur* because of limited acorn production in some regions. Seeds from additional provenances were thus obtained from local collaboration partners forming standalone provenances (Italy, southern Switzerland, southern France and Serbia, see Figure 1). Due to this imbalance creating unequal detection likelihoods, we focused our analyses on species-specific large-scale climatic adaptation rather than parallel adaptation patterns between species and regional adaptation. However, local provenances were not combined into single regional provenances for our analysis due to known soil moisture differences. Acorns of *Q. petraea* were also collected and planted. However, these were excluded from analyses after species identification (see below) because of a limited sample size in southern European provenances.

**Figure 1.**
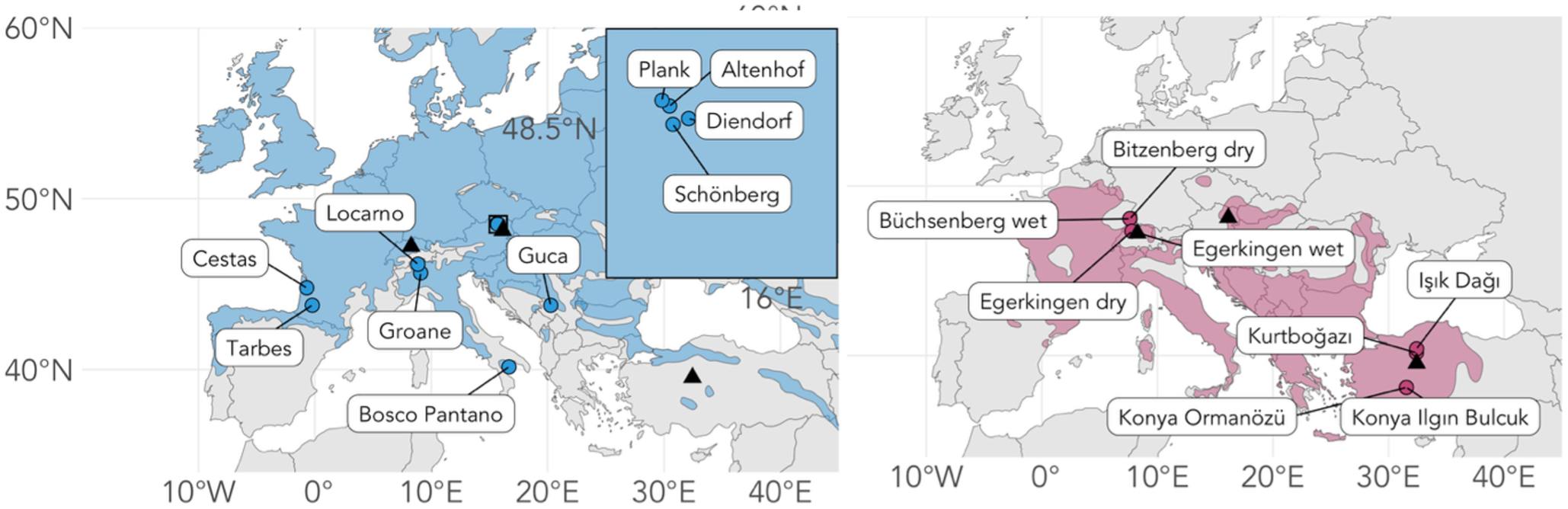
Locations of provenances (circles) and distribution ranges for Q. robur (left, blue) and Q. pubescens (right, red) grown in the three common gardens (triangles) located in Switzerland (near Zurich), Türkiye (Ankara), and Austria (Vienna). The box insert on the left shows the sampling locations within Austria.

At each site, 50-100 acorns were collected from at least six mother trees (Table S1). Acorns were collected either using ground nets or directly from the tree crown. Leaf or bud samples were taken from mother trees to confirm relatedness through molecular genomic methods (see below). After collection, acorns were treated with a thermotherapy in a water bath (41°C for two hours) to remove surface fungi and identify non-viable acorns through a float test. Acorns from Switzerland and Germany were collected in 2020 because it was a masting year, and thus were stored for 1 year at −2°C and 40-45% relative humidity in paper bags at Staatsklenge Nagold (Germany), to allow synchronized planting with the rest of the provenances that were collected in 2021.

Three common gardens were established in Türkiye (Ankara, 39.898°N, 32.743°E) Switzerland (Birmensdorf, near Zürich, 47.362°N, 8.455°E) and Austria (Vienna, 48.219°N, 16.240°E) in December 2021. At each location, the two species were planted in a fully randomised design within species blocks. To reduce competition between seedlings, acorn planting locations were spaced 30-45 cm apart. All acorns were planted directly into the garden without use of a nursery. We aimed for six seedlings from six mother trees per garden with the same mother replicated across all three common gardens. To protect against the high expected rate of germination failure (particularly from stored acorns), three acorns from the same mother were planted at each location and random surplus individuals were removed after germination in July of 2022 (year 1). To limit the loss of seedlings from predation, acorns were protected after planting with metal or plastic mesh. These protections were removed in the summer of year 1 to ensure they did not impede growth. Weeds and grass were removed before planting and all sites were weeded when necessary. All seedlings in Austria and Switzerland were treated against powdery mildew fungus in year 1 and year 2. Powdery mildew was not present in the experiment in Türkiye.

Due to the extreme drought in the summer of year 1 (2022), bare earth was covered at all three sites with straw or an anti-weed membrane. In the first growing season, drought related death cannot be easily distinguished from germination failure. Thus, sites were watered over the summer when needed in year 1 (weekly in Türkiye and every 2 weeks in Switzerland and Austria). In year 2, this was reduced to a minimum in Switzerland and Austria (an average of once every 3 weeks).

### Trait data collection

In both growing seasons, key growth and drought-related traits were measured for seedlings (Table S2). All measurements were conducted within a consistent 1-2 weeks period in each year to minimize temporal variation.

The total number of leaves per seedling was recorded at the end of August, after leaf expansion had ceased, but before senescence. All fully developed leaves present on each seedling were counted and those removed for sampling other traits earlier in the season (see below) were added to the total.

Seedling height measurements were conducted at the end of each growing season (November). Total height was measured to the nearest 0.5 cm using a metre rule positioned vertically at the base of the stem. Height was recorded as the distance from the stem base to the uppermost photosynthetic tissue. Diameter was measured 3 cm above the soil surface using digital callipers with millimetre precision.

Leaf traits were also recorded, these included: SLA, leaf dry matter content (LDMC), stable carbon isotope composition (stable carbon δ¹³C and nitrogen δ¹^5^N isotopes), leaf wet mass, and leaf dry mass. A single expanded, representative, mature leaf was collected from the sun-exposed outer canopy of each seedling and used for all traits. Leaves showing substantial herbivore or pathogen damage were avoided where possible. Leaves were collected during the morning (before 12:30) at the beginning of September in both growing years. Sampling was avoided during rainfall, and seedlings were not watered before collection. Immediately after collection, each leaf was wrapped in moist paper towel, sealed in a labelled plastic bag, and briefly stored in a dark insulated container before being transferred for processing.

Wet mass, leaf area determination, and transfer to the drying oven were completed on the day of collection. Leaves were removed from the plastic bag and gently blotted dry. Wet mass was then measured on a microbalance (precision ±0.25 mg). Each leaf was then scanned on a flatbed scanner with a ruler and sample identification label to enable leaf area estimation with the open-source software *ImageJ*. Following scanning, leaves were oven-dried at 60°C for 48 h. After this, they were reweighed using the same microbalance to determine dry mass. From these measurements, LDMC was calculated as the ratio of oven-dry mass to fresh mass, and SLA was calculated as leaf area divided by oven-dry mass.

Dried leaf samples were then transported in silica gel to the Swiss Federal Research Institute for Forest, Snow and Landscape Research WSL (Switzerland) for processing for isotope composition (δ¹³C and δ15N). Because δ¹³C and δ¹ N integrate physiological responses over longer periods, they were included as complementary traits to capture drought-related and nutritional variation not evident from growth measures alone. Approximately 1.0 ± 0.1 mg of dried leaf material was weighed into pre-tared tin capsules using a high-precision microbalance. Capsules were carefully sealed by compressing and rolling them into compact pellets using forceps. Each sealed capsule was reweighed to verify the final sample mass before being placed into a labelled sample tray for analysis. All work surfaces and tools were cleaned with ethanol between samples to minimize contamination. Each capsule was then placed in a deep well 96-well plate and processed at the Isotope Lab of WSL (Switzerland) by combustion in an elemental analyser (iso-Earth, Sercon, UK) coupled to an isotope-ratio mass-spectrometer (HS2022, Sercon, UK). Leaf δ^13^C values are the ratio of ¹³C to ¹²C relative to the international standard VPDB, and these values are impacted by the opening and closing of stomata in response to changes in water availability and are considered a metric of intrinsic water-use efficiency (Damesin et al., 1997; Farquhar et al., 1989; Roussel et al., 2009). Leaf composition in lignin versus non-lignin compounds and the large differences in leaf mass does not bias the isotopic signal in bulk leaf material within (Ghouil et al., 2024), therefore δ^13^C can also be compared across species of Mediterranean and temperate broad-leaf forest tree species. Leaf ^δ15^N values reflect the ratio of ¹^5^N to ¹^4^N and is related to plant growth and metabolism through variation in nitrogen acquisition and internal nitrogen cycling (Mateus et al., 2021).

Top soil was also taken at 10 randomly chosen points in each experiment in year 2, dried and processed for average δ¹³C, δ^15^N, percentage carbon and nitrogen (Table S3). As with leaf isotopes, these values are ratios of ¹³C to ¹²C and ¹^5^N to ¹^4^N respectively. Soil pH was measure at the beginning of the experiment and was similar across all sites (7.14-7.54).

### Low coverage whole-genome sequencing

Leaf samples for DNA extraction were taken early in year 1 (July 2022) when seedlings had >6 leaves in total. For those seedlings with less than six leaves, this was delayed till the end of August. To minimize damage, only one fully grown, unhardened leaf was cut from each seedling. Leaves were placed into paper teabags and then stored in silica gel until DNA extraction. Samples from the Austrian and Turkish common gardens were shipped and delivered in person by a researcher, respectively, for DNA extraction at WSL (Birmensdorf, Switzerland), with the exception of some Austrian mother trees, where DNA extraction was done at BOKU (Vienna, Austria) and then shipped to WSL.

At WSL, DNA was extracted from 10-30 mg dried tissue using a KingFisher96 System (Thermo Fisher Scientific, Waltham, USA) with an oak-specific sbeadex maxi plant kit (LGC Genomics, Berlin, Germany). A 1% agarose gel was used to assess DNA quality for the entirety of samples on the first three extraction plates, then a random subset of 10% on proceeding plates. The Austrian mother trees were extracted at BOKU using DNeasy Plant Mini Kit by Qiagen following the manufacturer’s protocol and then shipped to WSL. DNA quantity and quality were assessed with a NanoDrop spectrophotometer (Thermo Fisher Scientific, Waltham, USA) and (for a subset of the samples) with Quantus (Promega Corporation, Madison, USA). Extractions were then shipped to Cornell University’s Biotechnology Resource Center for Qubit quantification, followed by skim Nextera library prep, and MiSeq guided pooling of libraries. Final libraries were sequenced on two full Illumina NovaSeq6000 runs (paired end 151bp) at Weill Cornell Medicine with a target sequencing depth of 1-6x. 6 samples (2 per garden) were extracted and sequenced twice as full technical replicate samples, leading to a total of 946 samples sequenced. Plate negatives were also sequenced (n=14) to ensure no contamination occurred. An additional sample set of seedlings (n=345) from central Europe initially planted at WSL in a nursery in 2020 for a separate experiment *(Q. robur; Q. pubescens; Q. petraea*) and 27 *Q. petraea* from southern Europe (both mother trees and seedlings) planted in 2021, were also sequenced and used for the species assignment analysis. This was particularly important for ensuring we had as locally representative *Q. petraea* reference populations for species assignment.

### Sequencing data alignment

Sequencing reads were first demultiplexed and mapped to the chromosome contigs of the *Q. robur* reference genome (version: PM1N; Plomion et al. 2018; Plomion C et al. 2016) with plastid sequences from the reference genome of Darwin Tree of Life (dhQueRobu3.1, GCF_932294415.1). Reads were directly mapped using BWA-mem2 with default settings (Li, 2013). After mapping, individuals with a mean coverage of <1 were removed (n=28; 16 mother trees and 12 seedling individuals) for the entire analysis.

### Species assignment

Mother tree species assignment was initially done visually using leaf morphological traits in the field. However, it is notoriously difficult to separate oak species from the section *Quercus* in the field due to the large phenotypic plasticity in the traits used for identification and due to extensive hybridization (Kremer et al., 2002; Rellstab, Bühler, et al., 2016; Reutimann et al., 2020). Regular hybridisation also makes it possible that seedlings are of mixed origin even if the mother tree was correctly identified. Therefore, it was necessary to confirm species assignments for all mother trees and seedlings with our genetic data before any analysis. To this end, an inter-species set of GLs were called for all three species with *bcftools* view (Danecek et al., 2021; Li, 2011: minimum allele frequency 0.01, two minimum/maximum alleles, SNP mutations, QUAL > 30, average depth between 2 and 10 reads, chromosome level contigs only). The inter-species GL were filtered using *vcftools* varFilter (Danecek et al., 2011) with default settings resulting in 21,118,931 GLs. To remove linked sites and ensure reasonable computational handling times, a random subset of 2 million GLs was selected using the Bash ‘shuf’ command. Putative hybrids were then identified using *ngsAdmix* (Skotte et al., 2013), testing K from 2 to 12, with three replicates runs for each value of K (Figure S1) and *CLUMPAK* was used for visualisation and averaging across replicate runs (Kopelman et al., 2015). Incorrect species assignments were clearly identifiable at K=3 where three species could be resolved, and individuals were removed when they belonged to the wrong species (ancestry >90%; 11 individuals). Hybrids (ancestry of > 40% belonging to two species-specific clusters) were more challenging to identify and were only clearly present in individuals assumed to be *Q. pubescens* (none were detected in *Q. robur*). *Q. pubescens* hybrid individuals were removed based on assignments at K=7. A K of 7 was chosen because it was where central and southern European *Q. pubescens* clearly separated into units that matched biological expectations. Individuals were excluded if they had ancestry of > 40% belonging to more than one *ngsAdmix* inferred group (n=8). Finally, all *Q. petraea* were removed from further analyses due to limited sample size.

### Intraspecific genotype likelihoods

To obtain intraspecific (i.e. species-specific) variant calls, the cleaned sample sets were rerun per species in *ANGSD* (Korneliussen et al., 2014): a lcWGS pipeline that provides GLs rather than hard calling SNPs, using the built-in *GATK* model and a beagle file output. To ensure alignments were of high quality, only reads with a mapping quality above 30 (minMapQ), uniquely mapped (uniqueOnly 1), properly paired reads (only_proper_pairs 1), and with ‘no bad’ filters were used (remove_bads 1). Bam headers were also compared across paired files to ensure they were compatible (checkBamHeaders 1). A C value of 50 was applied to adjust for excessive mismatches (C 50) and the per-base alignment quality flag (baq 1) was used (Korneliussen et al., 2014). To remove low depth regions with limited information, and erroneously high depth regions that could be repetitive regions, additional filters for minimum depth and sample number (half the total sample number per species) and a maximum depth (three times the standard deviation of the mean) were used (minInd, setMinDepth, setMaxDepth). GL calls filters were applied to remove low-quality sites using a minimum minor allele frequency of 0.025 (minMaf), a SNP p-value of 1e^-6^ (SNP_pval), a minimum quality score of 20 (minQ), and skipping tri-allelic alleles (skipTri-allelic 1). Finally, both major and minor sites were called (doMajorMinor 1).

To estimate a window size for linkage disequilibrium (“LD”) pruning, r^2^ values were calculated with *ngsLD* across the inter-species GLs (Fox et al., 2019), as well as 54 unrelated *Q. robur* individuals (one representative per family) from the intraspecies GLs. To minimize computational demands, LD was calculated for Chromosome 1 only. The distance needed to decline to <0.2 was found to be ∼100bp (Figure S2). A linkage-pruned set of GL calls was then obtained using each species list of GL calls and *vcftools* ‘thin’ command, which keeps the first site in a given window size (100bp). *ANGSD* was then rerun as above with the linkage-pruned site list using the “sites” flag to generate LD-pruned GL calls.

### Quantitative genomic analyses

Trait heritability was estimated for each species separately using *MCMCglmm* (Hadfield, 2010) and an animal model using a relatedness matrix estimated from the lcWGS data. To this end, genome-wide relatedness was estimated in *ANGSD’s NGSRelate* (Korneliussen & Moltke, 2015) on the linkage-pruned GL dataset. Runs of *NGSRelate* were divided across chromosomes to allow for parallelisation and then the average genome-wide pairwise relatedness was calculated. *NGSRelate* calculates 21 relatedness statistics. The estimator “KING” was chosen for our relatedness matrix because it does not rely on allele frequencies, making it more robust to the structure present within our dataset. It also uses identity-by-state allele sharing patterns that capture the shared alleles across the genome relevant for a *GWAS* (Manichaikul et al., 2010; Waples et al., 2019). KING estimates from *NGSRelate* were converted into a matrix for *MCMCglmm* by generating and solving a symmetrical matrix in R, then taking the nearest positive definite matrix.

Mother identity was confirmed using the KING relatedness estimators. Mismatches (i.e. individuals with low relatedness to the assumed mother tree) were identified by converting the pedigree into a matrix and using the function “match.G2A” to compare this with the KING matrix. Individuals where the difference between the pedigree and molecular relatedness were >0.25 were considered unrelated and their mother given a new ‘dummy’ identifier.

To ensure comparability with the GWAS analyses (below) the trait data included were cleaned before analysis to remove outliers and, if needed, transformed to meet assumptions of a normal distribution. All traits were also normalized to ensure GWAS effect sizes were comparable and thus normalized values were also used to calculate heritability. Traits were modelled as a Gaussian response.

The *MCMCglmm* model included as fixed effects common garden and experimental year. As well as the climate conditions at the seed origin (provenance), including annual temperature (CHELSA bio1, Karger et al. 2023), yearly precipitation sum (CHELSA bio12) and precipitation variability (CHELSA bio15). These three climatic variables were chosen to capture different components of drought while minimising multicollinearity across predictors.

Additive genetic effects were modelled using animal (i.e. the individual ID) as a random effect and an inverse genomic relationship matrix (GRM). Dam identity (i.e. mother tree ID) was included as an additional random effect to account for shared maternal effects. To allow additive genetic variance and residual variance to differ between years, both the animal and residual terms were fitted with unstructured covariance matrices across years (“us(year):animal and us(year):units”), permitting estimation of year-specific variances and genetic covariances between years. The models were run using the strong priors R and G1: V= diag(2), nu=2+0.002; G2: V =1, nu =1, alpha.mu =0, alpha.V=1000; with 750,000 iterations, a burn-in of 10,000 iterations, and a 100 iterations thinning interval. Models were validated according to the recommendations in de Villemereuil (2024), thus all traces were evaluated by eye to ensure proper mixing, and effect size calculation (ensuring all were substantially above 1000).

Trait heritability was calculated according to de Villemereuil (2024) with the inclusion of fixed effects and maternal variance. Additive genetic variance (V_A_), maternal variance (V_M_), and residual variance (V_R_) were extracted from the posterior distribution of each model (i.e. VCV). V_F_ is the variance explained by the fixed effects, estimated as the variance of the linear predictor (Var(*Xβ*)). Phenotypic variance (V_P_) was then calculated as

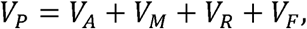

Narrow-sense heritability was calculated as standard

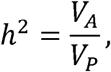

and the proportion of phenotypic variance arising from maternal effects was also calculated as

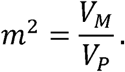

Estimates were calculated for each trait from every posterior sample and summarized using the mean and 95% credible intervals.

### Genome-wide association study (GWAS)

A GWAS was conducted to identify putative causative genes or genomic regions controlling trait variation. This was done separately for *Q. robur* and *Q. pubescens* using ANGSD *doAsso* with the latent genotype model (model 4; Jørsboe & Albrechtsen, 2022; Korneliussen et al., 2014). This model was chosen because it retains significantly more genotype information relative to other *doAsso* models. For both species, two binary covariates were used to account for the three common garden locations, as well as a species-specific number of covariates to account for relatedness within the dataset. Relatedness covariates were obtained by running *PCAngsd* with the linkage-pruned GL calls and transforming the covariance matrix output into eigenvalues (*eigen* in R v.4.3.1). The number of axes included in each GWAS was then established through testing, starting with the number corresponding to the levelling off point of each species’ PCA barplot and changing the number iteratively until test runs of *doAsso* (Jørsboe & Albrechtsen, 2022; Korneliussen et al., 2014) showed no *p-value* inflation (qqplots, package *gaston* Perdry et al., 2018). The first seven axes were used for *Q. robur* and the first two for *Q. pubescens*. The same number of covariates was used for all traits for a species. The minimum proportion of credible genotypes needed for a site to be included in the GWAS was set to 5% of samples.

Qqplots were examined by eye using the package gaston after running *doAsso*. If qqplots showed poor linearity, we retransformed traits (e.g. square-root transformation) and reran models. The model with the most linear qqplot was reported. Significant GWAS hits were determined for each trait using the Benjamini & Hochberg p values to control for multiple testing (Benjamini & Hochberg, 1995). P values were transformed using the R function *p.adjust* with “BH” as the method then the significance threshold of 0.05 was used on the adjusted values. As many solitary hits were identified, we also report those traits with clustered sites where more than one significant association was found within a 3000bp window (Coq--Etchegaray et al. 2023).

To identify putative site impact and gene function, associations with p< 5×10^-7^ were annotated in *SNPeff* (Cingolani et al., 2012). A fixed threshold was chosen because this is considered the ‘suggestive level’ in many GWAS (Chen et al., 2021). This allowed us to balance accuracy and information when contextualising the associations and ensured our findings can have broader relevance. To maximise the information obtained, sites were also mapped to the Darwin Tree of Life (“DTOL”) *Q. robur* reference genome assembly. It is a putative F2 hybrid with *Q. petraea* (*Mark Blaxter pers. comm.*) and thus was not used for our original mapping but is more comprehensively annotated than PM1N. To map the species to the genome, 500bp up- and down-stream of each site was cut from the PM1N reference genome with *bedtools* and then mapped to the DTOL genome with bwa-mem2 (default parameters). The re-mapped location of each site was then annotated using *SNPeff* and the resulting gene names extracted. For clustered hits, regional function was inferred using *genename* and *uniprot* (Seal et al., 2026; The UniProt Consortium, 2025).

## Results

### Genomic data

A total of 890 of 1322 sequenced samples were in our final data set. This was after removal of hybrid individuals (n=8) or species miss-assignments (n=27), *Q. petraea* references (n=10), samples with less than 1x coverage (n=28), sample negatives (n=14), and those from a parallel experiment used to aid species assignment (n=345). From these 890, 609 were *Q. robur* seedlings, 197 *Q. pubescens* seedlings, 80 individuals were mother trees, and 6 were full technical replicates (i.e. independent extractions and libraries from the same original sample). In *Q. robur* we identified 4,490,659 species-specific GL calls. In *Q. pubescens* we identified 4,894,041 species-specific GL calls. All 14 plate negatives were clean and contained no meaningful data.

Across the technical replicates, the mean KING values for both species were 0.45. The PCA and relatedness plots showing the underlying genetic and familial structure of the data is shown in Figure 2. High structure was found between southern and central populations of *Q. pubescens.* Within *Q. robur* the southern Italian provenance Bosco Pantano was highly distinct, followed by the two French provenances.

**Figure 2.**
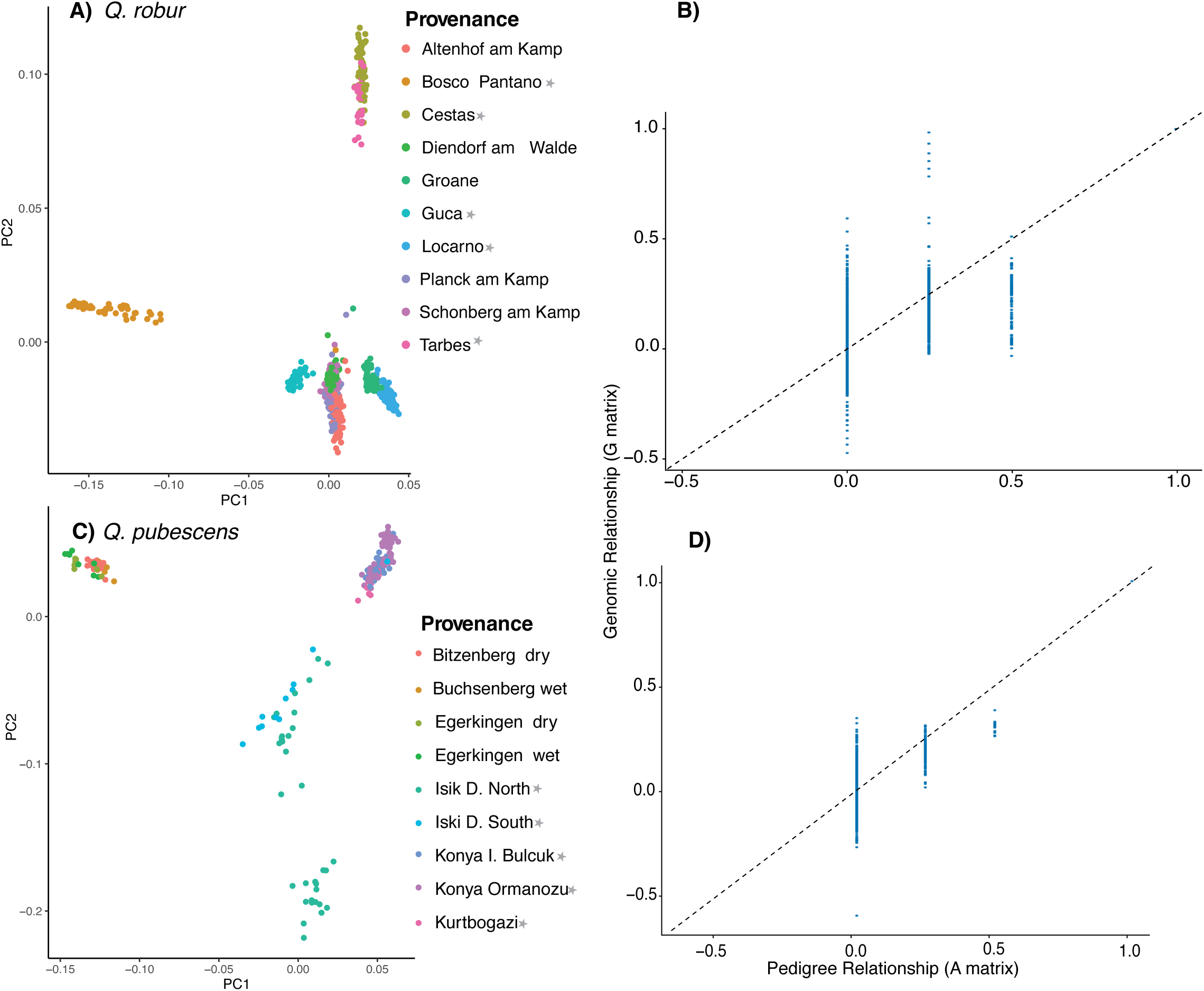
Principal components analysis of Q. robur (A) and Q. pubescens (C) calculated including half-siblings to show the relatedness structure of the data. The correlation between relatedness measured by KING and the uncorrected social pedigree (i.e. mother, half siblings) is shown in B) for Q. robur and D) for Q. pubescens. Grey stars represent Southern European provenances.

### Quantitative genomic analyses

All trait values are summarised across years and gardens in supplementary Figures S3 and S4, and Tables S4 and S5. *Q. robur* had noticeably larger leaves on average and was taller. The smallest trees (in height and many other growth-related traits) and subsequently lowest growth rate was in Türkiye. Values of δ^13^C were lower in *Q. pubescens*.

The narrow sense heritability of the measured traits was moderate to high in both species (Figure 3) and was often higher in year 1. Markedly similar levels of heritability were seen for many traits across the two species, including in key productivity traits like height and diameter in the second growing season. After accounting for relatedness, year and common garden frequently showed significant effects on many measured traits (Figure 3). Environmental variables from the provenance of origin were much less likely to have an effect, though notably impacted some leaf related traits in both species and growth-related traits in *Q. robur* (Figure 3). Maternal effects were also similar across the two species, but often higher in year 1 (Figure S5).

**Figure 3.**
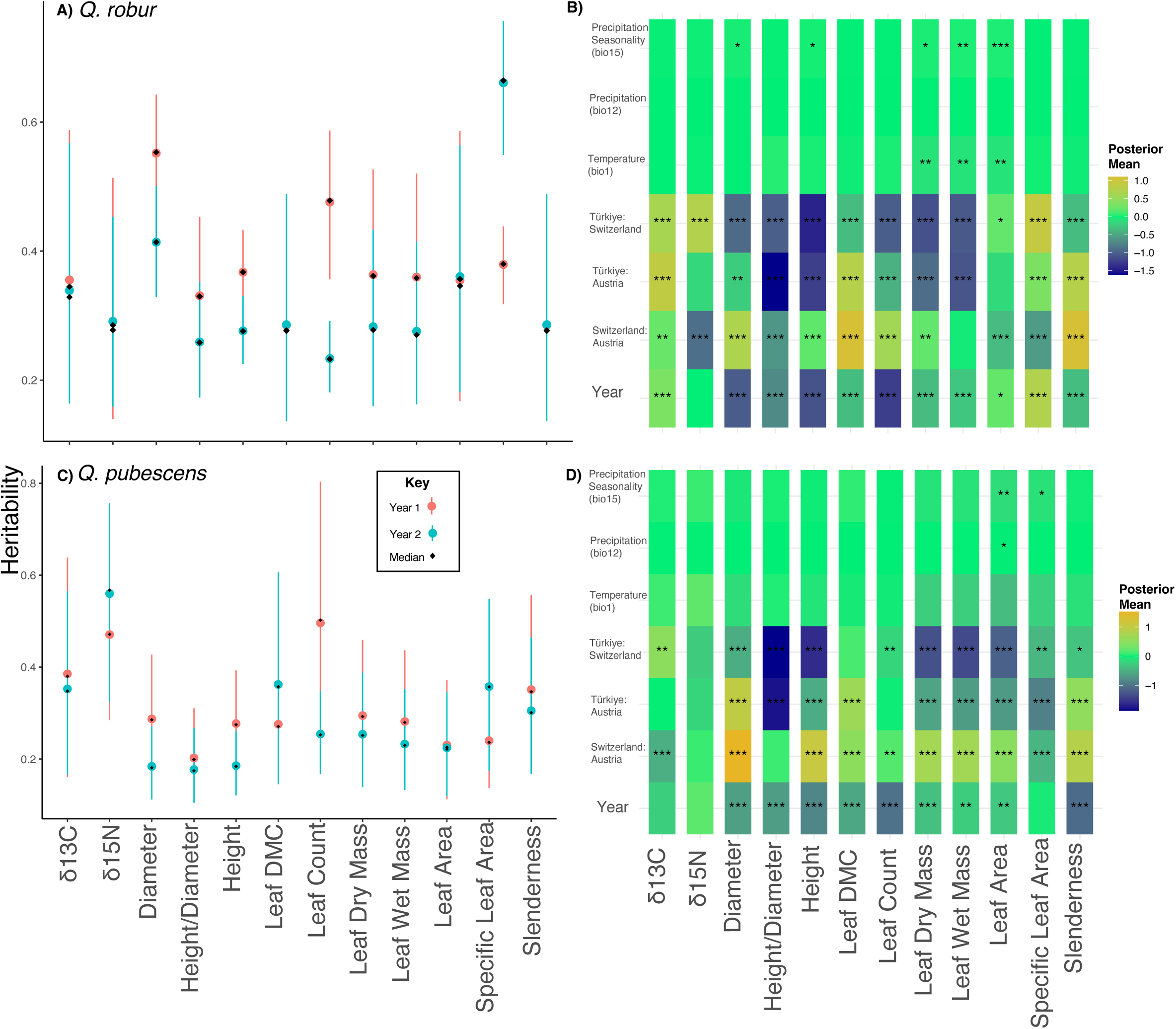
The narrow sense heritability of seedling traits for Q. robur (A) and Q. pubescens (C) calculated including fixed effects in MCMCglmm. The significance of fixed effects within the models are shown on the right (asterisks, Q. robur B; Q. pubescens D) with the magnitude of posterior mean effects sizes (colour intensity).

Looking at the posterior means in Figure 3, *Q. robur* seedlings from sites with higher precipitation seasonality were significantly (albeit marginally) taller, had thicker stems, as well as heavier and larger leaves. Similarly, in *Q. pubescens,* seedlings from drier sites with higher precipitation seasonality had marginally smaller leaves. In *Q. robur,* provenance temperature also showed a relationship with leaf mass and area, with seedlings from warmer sites having smaller and lighter leaves.

### Genome-wide association study

Table 1 shows the summary of GL (i.e. variant sites) significantly associated with traits in each year for each species (detailed breakdown in Table S6 and S7). 350 significant associations were found in *Q. robur* and 333 in *Q. pubescens.* <u>Q</u>qplots generally met expectations; however, two traits in *Q. pubescens* (growth rate height and LDMC) showed p-value inflation combined with deviations from the trait expectations of normality even after transformation (see Figure S6 and S7). These associations contain a number of putative associations with some minor of clustering (i.e. multiple GL that were close together and associated with a single trait) but because of the model violations we did not consider these results reliable (Manhattan plots in Figure S8 and S9). Without these traits, the significant associations *Q. pubescens* included 19 sites.

**Table 1.** The number of GLs (i.e. variant sites) significantly associated with traits in each year for each species. Shown are the loci with a p value of less than 0.05 after Benjamini & Hochberg p value conversion in R. In brackets are the number of sites considered “clustered”, i.e. where the number of other significant sites found within a 3000 bp window. Those with a * are traits that did not meet model assumptions of trait normality after transforation.

| Trait | Year | Number of sites associated<br>in *Q. robur* | Number of sites associated<br>in *Q. pubescens* |
| --- | --- | --- | --- |
| $\delta^{13}\text{C}$ | Year 1 (2022) | 0 | 0 |
| $\delta^{13}\text{C}$ | Year 2 (2023) | 206 (176) | 0 |
| $\delta^{13}\text{N}$ | Year 1 | 1 | 0 |
| $\delta^{13}\text{N}$ | Year 2 | 0 | 0 |
| Growth rate diameter | Year 1 & 2 | 0 | 1 |
| Growth rate height | Year 1 & 2 | 0 | 69 (2*) |
| Height/diameter ratio | Year 1 | 1 | 1 |
| Height/diameter ratio | Year 2 | 0 | 0 |
| Leaf dry mass | Year 1 | 5 | 0 |
| Leaf dry mass | Year 2 | 0 | 0 |
| Leaf dry matter content | Year 1 | 0 | 245 (11*) |
| Leaf dry matter content | Year 2 | 0 | 4 |
| Leaf surface area | Year 1 | 19 | 0 |
| Leaf surface area | Year 2 | 0 | 0 |
| Leaf wet mass | Year 1 | 47 (5) | 5 |
| Leaf wet mass | Year 2 | 1 | 0 |
| Seedling diameter | Year 1 | 11 | 0 |
| Seedling diameter | Year 2 | 1 | 1 |
| Seedling height | Year 1 | 5 | 1 |
| Seedling height | Year 2 | 0 | 0 |
| Slenderness (leaf count/height) | Year 1 | 1 | 6 |
| Slenderness (leaf count/height) | Year 2 | 6 | 0 |
| Specific leaf area | Year 1 | 39 | 0 |
| Specific leaf area | Year 2 | 7 | 0 |

For *Q. robur,* two traits showed clustered GWAS loci where more than one significant genotype (GL) association was found within a 3,000 bp sliding window. This was for δ^13^C collected in the year 2 (qqplot in Figure S10) and leaf wet mass in year 1 (qqplot in Figure S11). For leaf δ^13^C, a significant association spanning over 150kbp was identified on chromosome 11 (Figure 4A) within the cyclic nucleotide-gated ion channel 1-like exon. For leaf wet mass, two smaller association clusters were identified on chromosome 1 within the ethylene-responsive transcription factor ERF119 and on chromosome 6 within the vacuolar protein sorting-associated protein 35A-like exon (Figure 4B). Several solitary GL sites were associated with other traits or growing sea sons (Table 1).

**Figure 4.**
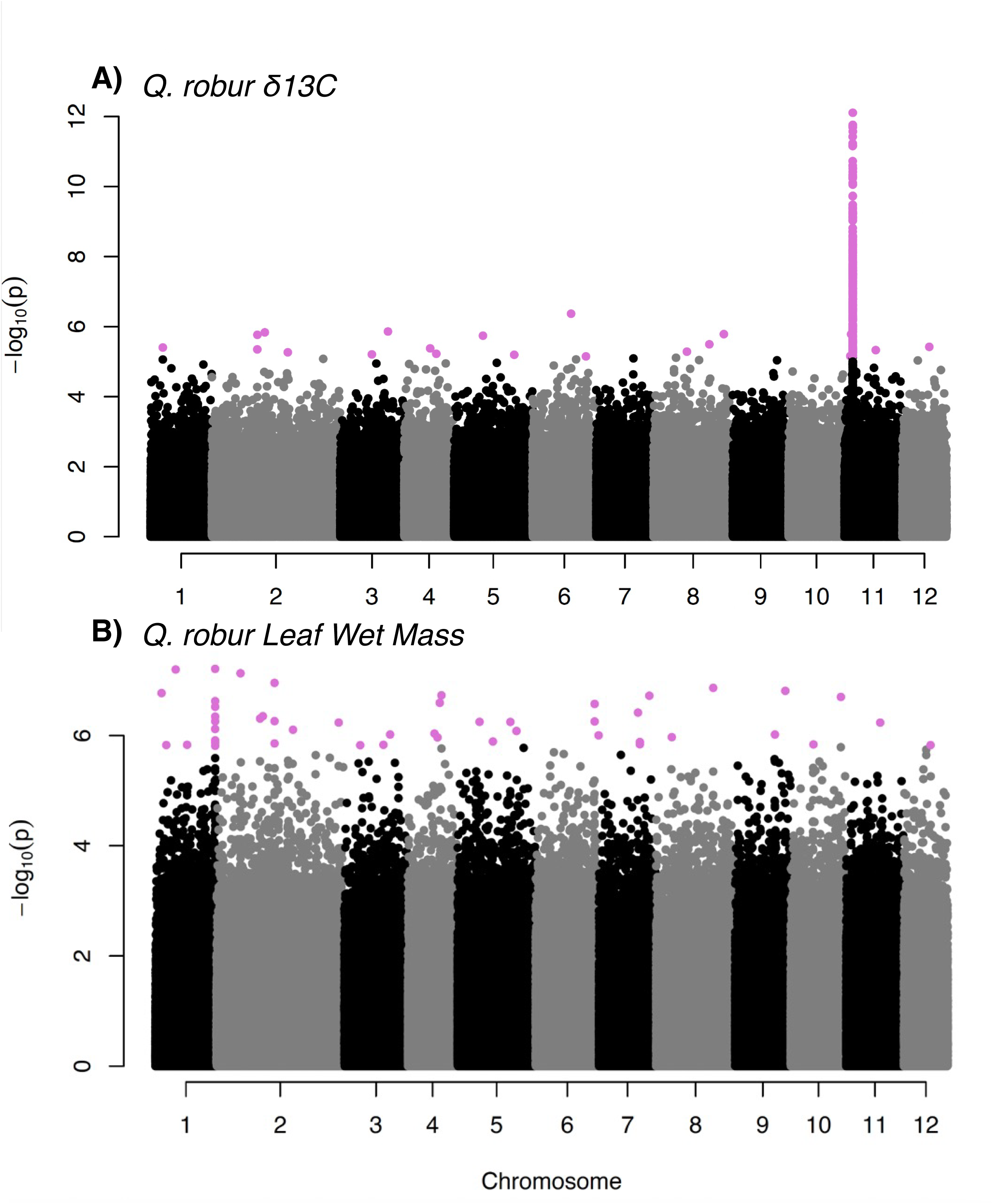
A) Significant GWAS hits across the genome in Q. robur for δ^13^C in year 2 of the experiment (top). B) Significant GWAS hits in Q. robur for leaf wet mass in year 1.

## Discussion

In this study, we combined common gardens of *Q. robur* and *Q. pubescens* with lcWGS to better understand climatic adaptation, as well as the genomic architecture of key traits in seedlings’ first two growing seasons, leveraging this information to inform assisted gene flow in both species.

Despite the use of GLs in place of SNPs, our relatedness values for technical replicates (0.45 i.e. close to the theoretical maximum of 0.5 for KING; Manichaikul et al., 2010; Waples et al., 2019) signalled high reproducibility in the dataset. Population structure also mirrored geographical distance between populations, likely reflecting isolation by distance patterns. Notably, the high distinctiveness of the *Q. robur* provenance Bosco Pantano met our expectations as this is an isolated relic of a distinct evolutionary significant unit within the species (Avanzi et al., 2025).

### Trait heritability and the impact of site of origin

Trait heritability was often high and values were generally similar across the two *Quercus* species, signalling substantial genetic control of early growth traits. Due to their economic importance, seedling height and diameter heritability have been widely quantified in tree species and thus offer the most context and insightful comparisons of the traits measured. In this study, the mean heritability of diameter was 0.55 and 0.41 for *Q. robur* (year 1 and 2, resp.) and 0.28 and 0.18 for *Q. pubescens* (year 1 and 2, resp.). For *Q. robur* these heritability values are well within the range previously reported under similar growth conditions; values of 0.47 were found in 20-year-old trees from a Danish common garden (Lobo et al., 2025), and values of 0.39-0.46 for 10-year-old trees grown in central European common garden (George et al., 2020). Though we are not aware of studies reporting diameter heritability values for *Q. pubescens,* values similar to both species’ have been reported for seedlings from other tree species. Focusing on studies examining seedlings of a similar age, this included 5-year-old seedlings of *Pinus koraiensis* (0.35-0.4; Lee, Oh, and Kim 2024), three-year-old *Eucalyptus spp.* (0.53-0.55; Resende et al. 2012), and two-year-old *Hevea brasiliensis* (0.217; Silva et al. 2014). These species span a large taxonomic range across four distantly related genera, suggesting that early life diameter is generally under substantial genetic control in trees, likely due to its close link with growth strategy, crown competition, and drought susceptibility (Jensen & Löf, 2017; Vander Mijnsbrugge et al., 2020).

Height heritability was moderate to high in our seedlings (0.36 and 0.27, *Q. robur* in year 1 and year 2, resp.; 0.27 and 0.18 *Q. pubescens* year 1 and year 2, resp.). These values are lower than previously reported values for subadult *Q. robur* (0.37 in 14-year-olds; Caignard et al. 2024; and 0.39-0.52 in 10-year-olds; George et al. 2020). However, this is likely an age effect because height heritability has previously been shown to change with age in young *Q. robur*, increasing in the first ten years of life (Bogdan et al., 2017). Similar changes with age in height heritability have been reported for other tree species (e.g. *P. koraiensis*; Lee et al. 2024; *Pinus taeda,* Xiang, Li, and Isik 2003; *Pinus pinaster,* Danjon 1994), collectively suggesting that early growth is under increasing genetic control as maternal effects decline. Once trees reached maturity, heritability often declines or levels off, likely because of the increasing importance of plastic adjustment to local conditions and competition (Stimm et al., 2021; Xiang et al., 2003).

The substantial heritability of virtually all the measured traits indicates early oak growth is under significant genetic control. While this provides a foundation for selection to act and generate local adaptation, many traits appeared to be also substantially plastic because of the persistent and strong impact of common garden signalling local adjustment to the environment (similar to that seen in *Q. petraea*, Sáenz-Romero et al. 2017). Accordingly, only a few of the growth and leaf traits we measured showed a symptomatic association with climate at provenance origin in either species. Similar limited associations with climate have been identified in *Q. petraea,* where only budburst was correlated with provenance temperature, while maternal investment was a better predictor of seedling growth (Losch, et al., 2025).

While our focal species are known to be generally drought tolerant, drought seasonality and timing impact the resilience of oak growth. Specifically, spring droughts are harmful to white oaks because they coincide with earlywood vessel growth (Bose et al., 2021; Popa & Popa, 2026). Previous relationships between drought and growth have been recorded for northern European provenances (Bose et al., 2021). Though a lack of an association with annual precipitation found in this study may seem surprising in *Q. robur*, provenances from the southern edge and central core of *Q. robur*’s range, like those included here, have previously been found to show a similar lack of association between growth and annual precipitation (Bose et al., 2021). This suggests the associations are regional and may require comparisons with more northern provenances to be detected. Importantly, the lower δ^13^C in *Q. pubescens* signals consistent inter-species differences in carbon isotope discrimination and intrinsic water-use efficiency, supporting the higher general drought tolerance generally of *Q. pubescens* (Ghouil et al., 2024).

The signal of climatic adaptation in *Q. robur* leaf area is also supported by other studies. Leaf area is currently declining under selection from rising temperatures in *Q. robur* in France (Caignard et al., 2024). Similarly, leaf morphology has been shown to be highly adaptive between the temperate and Mediterranean species, with traits converging across climates rather than across the phylogeny (Martín-Sánchez et al., 2024). Leaf-related traits were thus investi-gated in greater depth in a follow up study (McNamara et al., *In prep*).

### GWAS of seedling growth and drought-related traits

The GWAS revealed associations with key drought and growth traits in both species, with three “classical” Manhattan plot peaks in *Q. robur*. Other associations were scattered, potentially reflecting the low linkage disequilibrium, gaps from lcWGS, or the polygenic nature of traits (Crouch & Bodmer, 2020).

The most striking GWAS result was the identification of a 150kbp genomic region on chromo-some 11 spanning an exon of the predicted cyclic nucleotide-gated ion channel 1 in *Q. robur,* which was associated with leaf δ^13^C in year 2. Validating this result, a previous quantitative trait loci (“QTL”) study on leaf δ^13^C in *Q. robur* identified multiple QTLs on chromosome 11, including two can be considered physically close to our result when accounting to the difference in marker resolution (Brendel et al., 2008). Notably, these QTLs were of major effect, estimated to account for over 20% of the trait phenotypic variance (Brendel et al., 2008). Leaf δ^13^C values are impacted by the opening and closing of stomata in response to changes in water availability and are considered a metric of intrinsic water-use efficiency (Farquhar et al., 1989). In *Arabidopsis thaliana,* stomata are controlled by the plant hormone abscisic acid (ABA), which triggers Ca^2+^ changes in the guard cells surrounding the opening leading to closure (Bauer et al., 2013; Tan et al., 2023). The gene identified in our GWAS is part of the cyclic nucleotide-gated channel (“CNGC”) gene family, which are the channels used for the ABA-activated Ca^2+^ movement into guard cells (Tan et al., 2023). *A. thaliana* knock-outs of CNGC5/6/9/12 have greatly reduced stomatal opening (Tan et al., 2023). Collectively, this indicates that the association between δ^13^C values and CNGC1 in *Q. robur* has likely arisen through the genes’ role in the stomatal closure signalling pathway. The CNGC1 gene family role in guard-cell Ca² signalling and stomatal regulation, provides a plausible mechanistic link to variation in δ¹³C. Leaves were sampled in September, thus δ^13^C values represent stomatal responses to drought across the growing season and this association is consistent with genetic variation in intrinsic water-use efficiency and drought-responsive carbon balance (Yang et al., 2026).

Interestingly, despite the strong relationship in year 2 and the same sampling protocol for δ^13^C in year 1, the association between trait values and the CNGC1 region was not identified in year 1, the first growing season. Though we made the best efforts to control conditions across the years, the lack of a repeated patterns across the two years may be because of phenotypic plasticity (different summer conditions), maternal effects (e.g. use of maternally supplied seed resources in year 1), or the effect of watering. All of these factors could have altered the strength of the δ¹³C signal or added noise that obscured the signal in year 1. Notably, experimental year 1 was in 2022, a particularly hot and severe drought year in central and southern Europe (Bevacqua et al., 2024). Though drought is precisely when stomatal closure will be extremely important to prevent water loss, the unequal germination dates (e.g. spanning late April to early August in the Swiss common garden) could have also obscured the genomic association by creating uneven growing season lengths captured by the δ^13^C values.

For *Q. robur* leaf wet mass, two smaller genomic regions were associated with the trait only in year 1. These were on two separate chromosomes, one within the vacuolar protein sorting-associated protein 35A-like exon (VPS35) and other in the ethylene-responsive transcription factor (ERF119). Vacuolar protein sorting genes, including VPS35A/B/C, have previously been shown to impact plant growth and leaf senescence in *A. thaliana* and this may drive part of the relationship observed here in oaks (Yamazaki et al., 2008). Importantly, VPS35 also plays a role in the storage of proteins and thus is expressed during *A. thaliana* embryogenesis, as well as during seed germination itself (Yamazaki et al., 2008). Previous explorations into early growth of oaks have shown the first growing season is heavily reliant on the maternally supplied seed resources (Leverkus et al. 2026; Losch, et al. 2025; Quero et al. 2007). Maternal effects were clearly visible in our experiment and above 10% for leaf wet mass. It is likely that the GWAS relationship we identified with leaf wet mass is also driven by VPS35’s role in early growth. Supporting this, a GL in the VPS35 exon also showed significant, albeit single unclustered associations with year 1 diameter, leaf area, and leaf dry mass, as well as year 2 diameter.

Leaf wet mass also showed repeated associations in the GWAS with ethylene-responsive transcription factor family genes (ERF) including a clustered association in ERF119, as well as individual GL associations with ERF120/121/122. The ERF gene family is composed of transcription factors that play key roles in ethylene signalling, stress responses and development (Müller & Munné-Bosch, 2015). Notably, ERF genes are involved in dehydration responses in *A. thaliana*, likely explaining the relation with wet mass (Sakuma et al., 2002). The top scoring ERF119 GL for the leaf wet mass GWAS also showed additional unclustered association with leaf area in year 1. Another GL in this gene is associated with diameter in year 2, suggesting a potential role of ERF119 in growth generally perhaps through an impact on ethylene signalling (Müller & Munné-Bosch, 2015).

Beyond the two clustered hits discussed above, additional significant associations were found at single, isolated, sites in both species. Even though they do not have the classical GWAS peaks, these associations should not be discounted as type 1 errors, due to the low linkage disequilibrium in white oaks that spans only a few hundred base pairs (Nocchi et al. 2022 and 100bp, this study). Supporting this, around 5% of significant GLs in *Q. robur* show associations with multiple traits, including highly co-dependent traits like leaf wet mass and leaf dry mass. Many of these GLs are also found within genes with plausible functions. For example, a GL in the UDP-arabinopyranose mutase 2 gene was associated with *Q. robur* leaf wet mass and dry mass in year 1 and is known to play a role in cell wall synthesis. Associated GLs are listed in the supplementary material as resource for future research (Table S6 and S7).

For *Q. pubescens,* fewer GLs were identified in the GWAS and there were no clustered hits. This likely reflects the smaller number of populations and narrower geographic range causing a loss of statistical power (M. Wang & Xu, 2019). Focusing on δ^13^C, 3 of the 4 significant associated GLs were in uncharacterised genes. The remaining annotated GL was in the intergenic region of a nerolidol synthase (NES) 1-like gene. NES is a plant enzyme that produces volatiles (linalool or nerolidol) used in internal signalling (Ashaari et al., 2021). Once stomata close the CO_2_ availability falls, changing reaction rates within the cell and this leads to the biosynthesis of secondary compounds like linalool or nerolidol (Kleinwächter et al., 2015). We speculate that the relationship with δ^13^C could in fact reflect their shared trigger rather than the genomic architecture of ^13^C metabolism itself. Further GWAS explorations into this species should be conducted due to its high drought tolerance. A species-specific reference genome and annotation would potentially support this by helping ensure function can be inferred.

### Results in light of assisted gene flow

*Q. robur* and *Q. pubescens* are thermophilic, drought-tolerant (particularly *Q. pubescens*), and ecologically important keystone species in many regions of Europe, making them central to maintaining forest function under climate change (Wöhlbrandt et al., 2026). The growing evidence of their broad drought tolerance and high genetic diversity across the southern and central parts of their range (Schotte et al., 2026; Niemczyk et al., 2025; Wang et al., 2025), signals limited need for widespread assisted gene flow programmes in these regions (though see productivity changes in Wöhlbrandt et al., 2026). Our results add to this, indicating that though climatic adaptation to seasonal precipitation variability is present in seedling traits in central and southern Europe (including in economically important growth traits), these relationships have limited impacts on trait values in the first two years of life. Nevertheless, southern white oak populations have recently experienced elevated drought-induced mortality (Bose et al. 2021) and, in some cases, are experiencing fast genetic erosion (Avanzi et al., 2025; Morcillo et al., 2020). Furthermore, in other studies, greater sensitivity in more northern provenances and signs of adaptation to precipitation have been recorded for white oaks (Gosling et al., 2024; Neumair et al., 2022; (George et al., 2020; Rellstab, Bühler, et al., 2016). These signal that the strategy of no-intervention is not universally suitable and carries the risk of local productivity loss and reduced forest health.

Targeted management interventions should be considered for European white oaks, especially into northern populations, highly drought stressed populations that show productivity loss, or genetically depleted populations. Interventions are necessary to help maintain generational succession (e.g. ensuring seedling establishment if adults are too stressed to produce acorns), ensure geneflow which will bolster local genetic diversity and thus adaptive adaptative potential, and to improve forest resilience long-term. Our results indicated that targeted interventions should use seeds and seedlings from provenances exhibiting local adaptation or exposure to high seasonal precipitation variability and with smaller leaf sizes, because this would allow managers to introduce climatic adaptations that we have shown impact seedlings but have also been shown to be important in both seedlings and adult trees (e.g. Caignard et al., 2024). Though oaks have high genetic diversity and thus adaptive capacity, the dramatic pace of climate change will likely overtake natural adaptive timescales and these actions will help oaks keep pace. Importantly, they are in keeping with previous recommendations of mixing based on studies of adult trees (Girard et al., 2022). Neither high seasonal precipitation variability nor leaf sizes is universally correlated with latitude; colder environments can also show high seasonal precipitation variability (Reichmuth et al., 2025) and provenances can have huge variation in leaf size (this study and Caignard et al., 2024). This offers the chance for smaller scale transfer with a reduced risk, legal barriers, and thus lower implementation costs. An alternative, minimalist, management approach would be to consider seasonal precipitation variability and leaf sizes when establishing new seed orchards or selecting mother trees for plantations. This would ensure maladapted seed material is not being used for management that is already ongoing. Both represent feasible and necessary strategies to help stressed white oaks stands keep pace with rate of contemporary climate change by offering fuel for adaptation.

## Conclusion

In conclusion, we found that seedling trait heritability was often high and similar in *Q. robur* and *Q. pubescens*. There was limited evidence for climate adaptation in seedling traits, with some signals of adaptation to precipitation seasonality (i.e. more irregular precipitation) and in leaf size. A large GWAS hit was identified on chromosome 11 in *Q. robur* that was associated with δ^13^C values suggesting there is genetic variation in intrinsic water-use efficiency present within populations. Managers interested in introducing climatic adaptations into stressed populations for climate smart forestry could consider these findings when searching for suitable seed sources for transfer material.

## Supplementary Material

Table S1: Provenance collection details and planting numbers

Table S2: Trait data collected for oak seedlings across all three common gardens in 2022 and 2023

Table S3) surface soil stable isotope analysis across ten random within plot locations for each garden

Table S4) trait data collected across the two years for Q. robur Table S5) trait data collected across the two years for Q. pubescens

Table S6) Those GLs exceeding the threshold of p< 5×10-7 for Q. robur. Shown on the right is the PM1N annocation and on the left the DTOL annotation.

Table S7) Those GLs exceeding the threshold of p< 5×10-7 for Q. pubescens. Shown on the right is the PM1N annocation and on the left the DTOL annotation.

Figure S1: Averaged ancestry coefficients estimated using ngsAdmix across three white oak species (*Q. petraea* included). Tested K=2-12, with three replicates runs for each value of K. In K=3, orange trees represent *Q. robur* and blue tree *Q. pubescens*.

Figure S2: Linkage disequilibrium between GL calls as estimated by ngsLD constrained to chromosome 1 for computational requirements

*Figure S3:* Q. robur trait value violin plots divided across the three gardens (x axis) and years (colours).

Figure S4: *Q. pubescens* trait value violin plots divided across the three gardens (x axis) and years (colours).

Figure S5: Maternal effect as estimated from MCMCglmm

Figure S6: qqplot for the Leaf Dry Matter Content in Year 1 for *Q. pubescens* from the doASSO GWAS analysis and gaston.

Figure S7: qqplot for the Height Growth rate in Year 2 for *Q. pubescens* from the doASSO GWAS analysis and gaston.

Figure S8: Manhatten plot for the Leaf Dry Matter Content in Year 1 for *Q. pubescens* from the doASSO GWAS analysis and ggmanh. Shown in coral are sites with significant p values after the Benjamini-Hochberg correction for multiple testing.

Figure S9: Manhatten plot for the Height Growth rate in Year 2 for *Q. pubescens* from the doASSO GWAS analysis and ggmanh. Shown in coral are sites with significant p values after the Benjamini-Hochberg correction for multiple testing.

Figure S10: qqplot for the δ^13^C in Year 2 for *Q. robur* from the doASSO GWAS analysis and gaston.

Figure S11: qqplot for the leaf wet mass in year 1 for *Q. robur* from the doASSO GWAS analysis and gaston.

## Authorship contributions

Data collection: Leigh DM, Acar P, Blyth C, Jansen SA, McNamara S, Vitali V, Kaya Z Data analysis: Leigh DM

Writing: Leigh DM, Acar P, Jansen SA, Neophytou C, Rellstab C, Blyth C, Popovic V, Kremer A, Piotti A, Graf R, McNamara NS, Vitali V, Saurer M, Kaya Z

Conceptual input: Saurer M, Kaya Z, Neophytou C, Rellstab C

Sampling: Leigh DM, Acar P, Graf R, Jansen SA, Jovanović S, Kaya Z, Kremer A, Mavi İdman O, Neophytou C, Piotti A, Rellstab C,

## Acknowledgements

With thanks to our DAAD Rise summer students for their help in the experiment: C. Gowenia, M. Shah, C. Altendorfer, M. Notter. Thanks to the BOKU interns. Thanks to S. Brodbeck, A. Braun, M. Castellaneta, A. Lapolla, O. Pericolo, M. Rosito, S. Pfister, R. Milčevičová, J. Stucki, M. Walser, M. Oettli, G. Golesch, A. Malaspina, A. Papadopoulou, Lazic S., G. Rabong, E. Zimm and S. Gruber for field work, logistical and contributions. Thanks to C. Grossiord, M. Schuman, and Y. Vitasse for their scientific and methodological input. Thanks to G. Reiss and the experimental garden team at WSL for their support. F. Reinthaler, F. Puchinger, D. Soukup, P. Fischer, are the garden team at Knödelhütte. Thanks to R. Köchli and J. Luster for soil sampling and analyses. Thanks to the Genetic Diversity Centre (GDC) of ETH Zurich (N. Zemp) for bioinformatic support. We thank K. Duyar, S. Halici and B. Altun from collection department/NBGT for their support in establishing and maintaining the Turkish common garden. Thanks to R. Kaufman, O. Bauer, F. Aravanopoulos, D. Semizer-Cuming, M. Oettli and M. Argelich Ninot. Thanks to T. Ebinger, Seed storage Nagold and Staatsklenge. The authors acknowledge the use of ChatGPT (OpenAI, GPT-5.5) to automate and troubleshoot iterative analyses of MCMCglmm model outputs and to create some paneled figures for visualization. All generated code was independently reviewed, stress tested manually on the study data by the lead author. The interpretation of results, diagnostic plots, methodological decisions, and scientific conclusions are solely those of the authors and were done by eye.

## Permit numbers and information

Compliance with National Access and Benefit-Sharing Requirements were managed by ACORN leads PA, CN, and CR: Türkiye is not a Party to the Nagoya Protocol; instead, access to and use of the Turkish plant material followed national regulatory procedures administered by the competent authorities of the Republic of Türkiye, and all associated documentation can be provided to the editor as Supplementary Files or under Data Availability. Depending on whether the material comprised seeds, leaf tissue, or derived materials (including DNA), it was subject to three distinct permission and documentation routes, all obtained from competent departments of the Ministry of Agriculture and Forestry of the Republic of Türkiye: (i) sampling permissions for seeds and leaf tissue were obtained from the General Directorate of Nature Conservation and National Parks (DKMP) (Ref. No: E-21264211-288.04; Attachments 1–2: (ii) research permit and cooperation protocol); the establishment of common garden trials with Turkish seed material was authorised through the Aegean Agricultural Research Institute / National Seed Gene Bank under an executed Material Transfer Agreement (MTA), and through the General Directorate of Forestry (Ref. No: E-36178555-604.99-2687270) in accordance with applicable Turkish forestry legislation, including the export permit, import permit, export pre-permit (Ref. No: E-59252610-435.01.01-2804607), import pre-permit (Ref. No: E-59252610-435.01.02-2804663), and phyto-sanitary certificate (No: EC/TR A 4636521) (Attachments 3–9: MTA, forestry permit, ex-port/import permits, export pre-permit, import pre-permit, and phytosanitary certificate, respectively), with the material subsequently cleared upon arrival through the corresponding Swiss phytosanitary import permit (No: CH 0010501; Attachment 10); and (iii) transfer of tissue and DNA for genomic analyses was processed through the General Directorate of Agricultural Research and Policies (TAGEM) (Ref. No: E-57776492-730.06.01-2784411; Attachment 11), together with the associated tissue transfer permit (Ref. No: E-94471571-903.07.03-6711640; At-tachment 12).

Austria, Germany, Serbia and Switzerland: are Nagoya members, but do not require Prior Inform Consent and Mutually Agreed Terms. The access is free. In France, which is also a member, access for research addressing silvicultural purposes are exempted from PIC and MAT (the answers of the focal points are saved). Italy is not a member of the Nagoya protocol. The focal points all confirmed that there are no ABS-related obligations.

