## Supplementary material for "Adaptive variation in drought-related traits across southern and central European white oak (*Quercus* sect. *Quercus*) populations": Figure S1

### CLUMPAK main pipeline - Job 1693833185 summary

#### Major modes for the uploaded data:

K=2

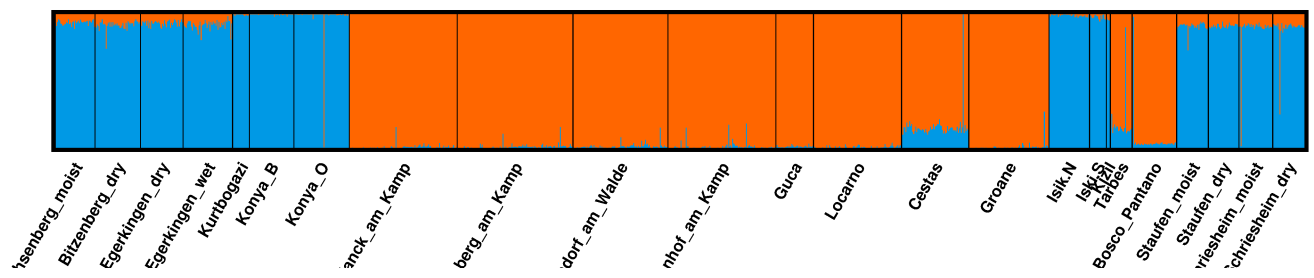


K=3

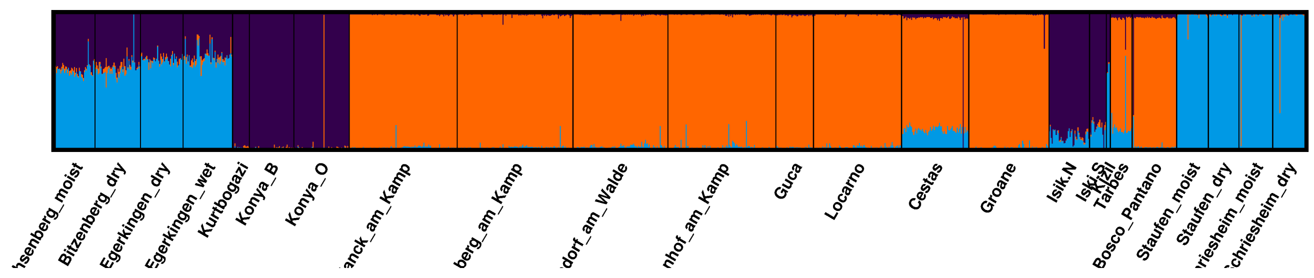


K=4

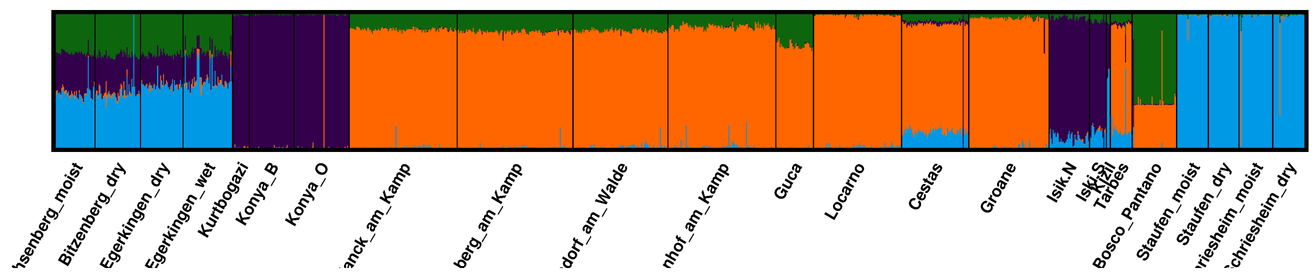


K=5

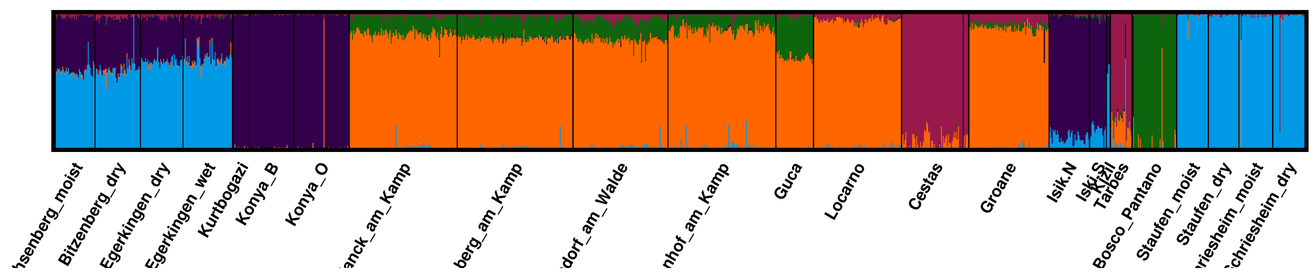


K=6

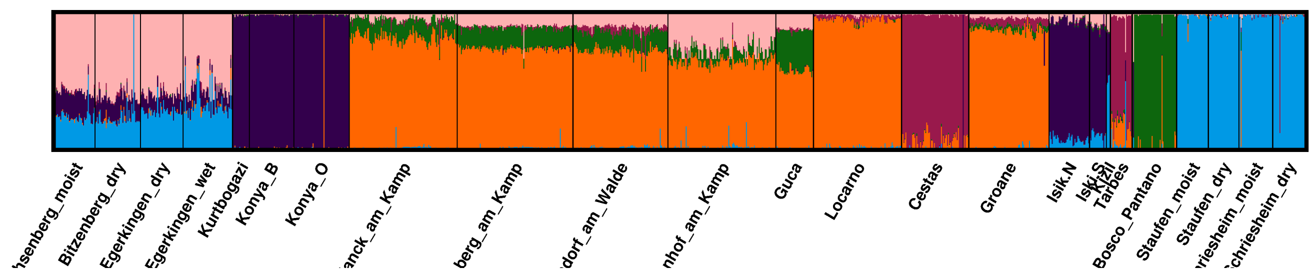


K=7

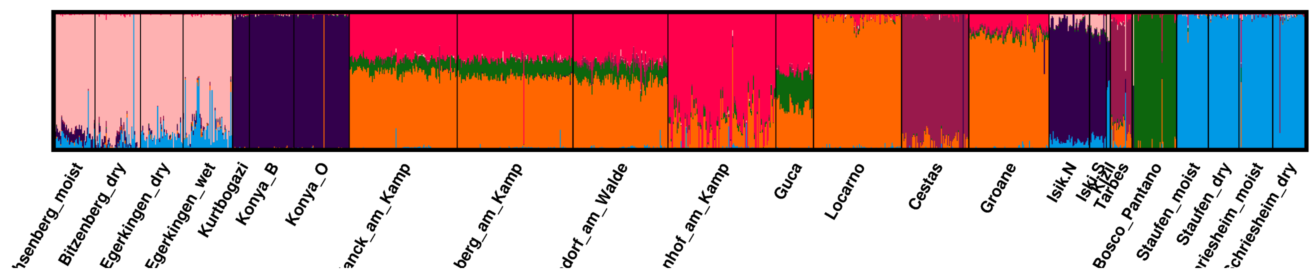


K=8

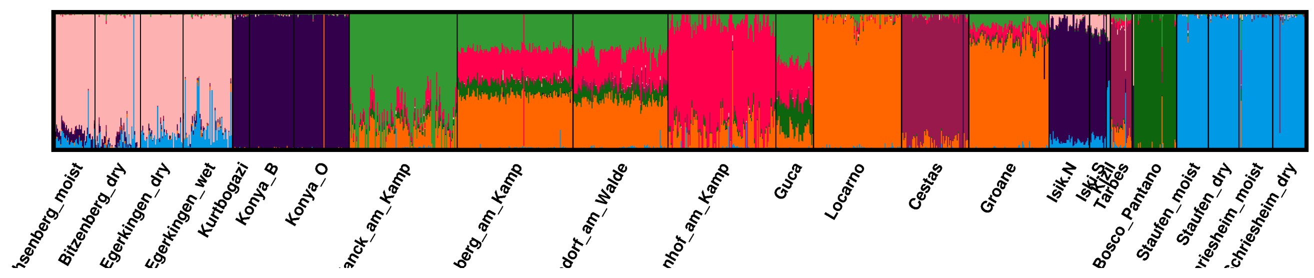


K=9

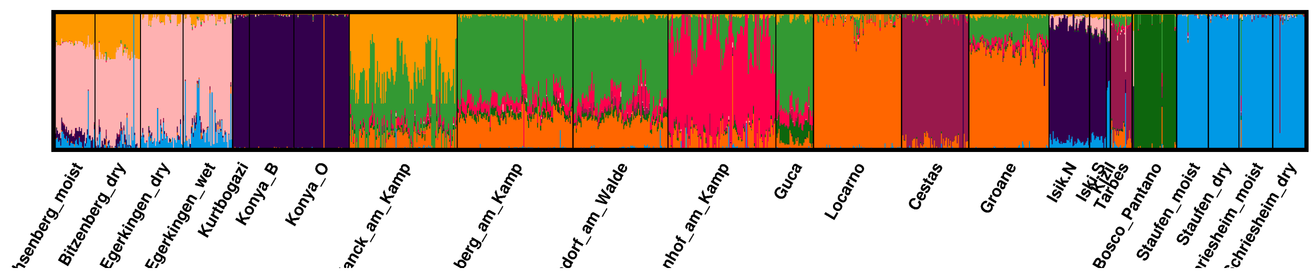


K=10

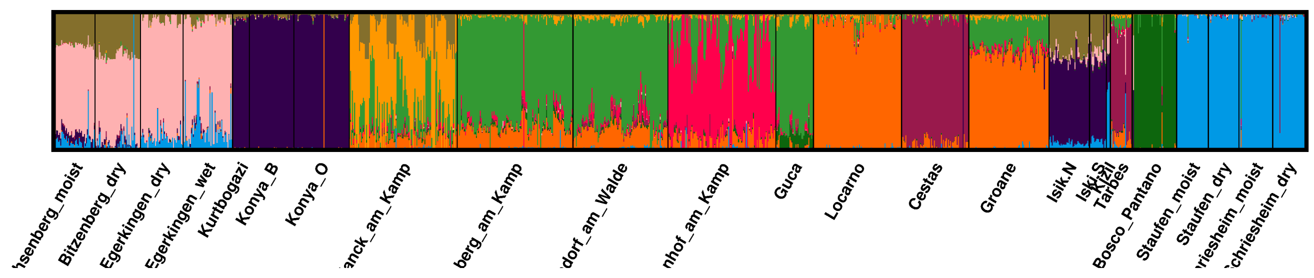


K=11

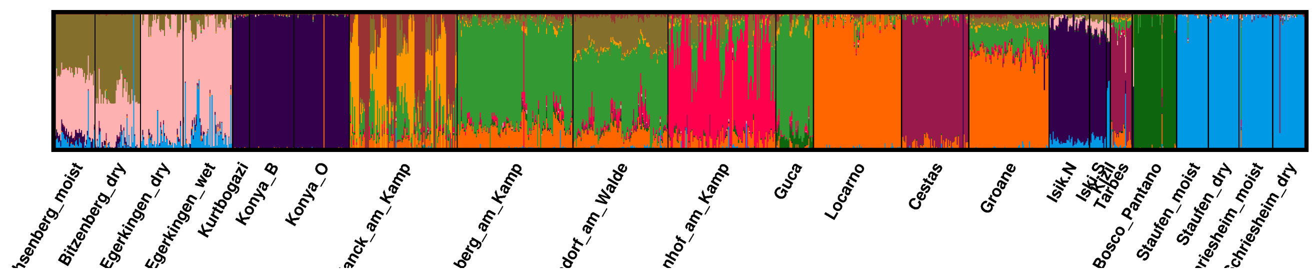


K=12

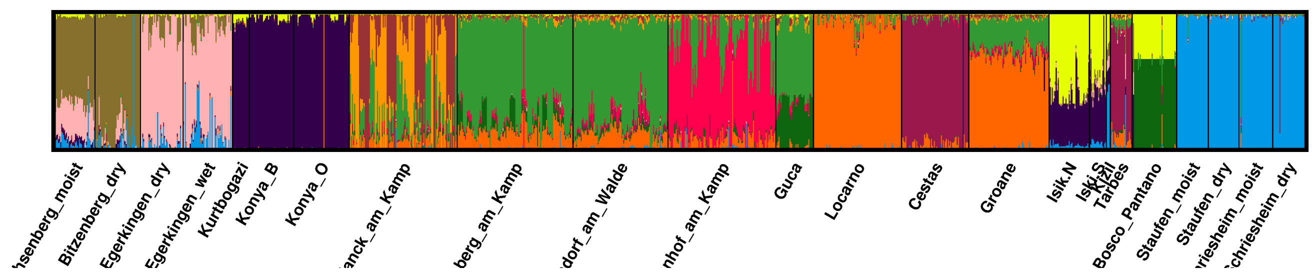


#### Minor modes for the uploaded data:

#### Division of runs by mode:

K=2 3/3
K=3 3/3
K=4 3/3
K=5 3/3
K=6 3/3
K=7 3/3
K=8 3/3
K=9 3/3
K=10 3/3
K=11 3/3
K=12 3/3
