## Supplementary figures and images for "Adaptive variation in drought-related traits across southern and central European white oak (*Quercus* sect. *Quercus*) populations"

### Figure S2

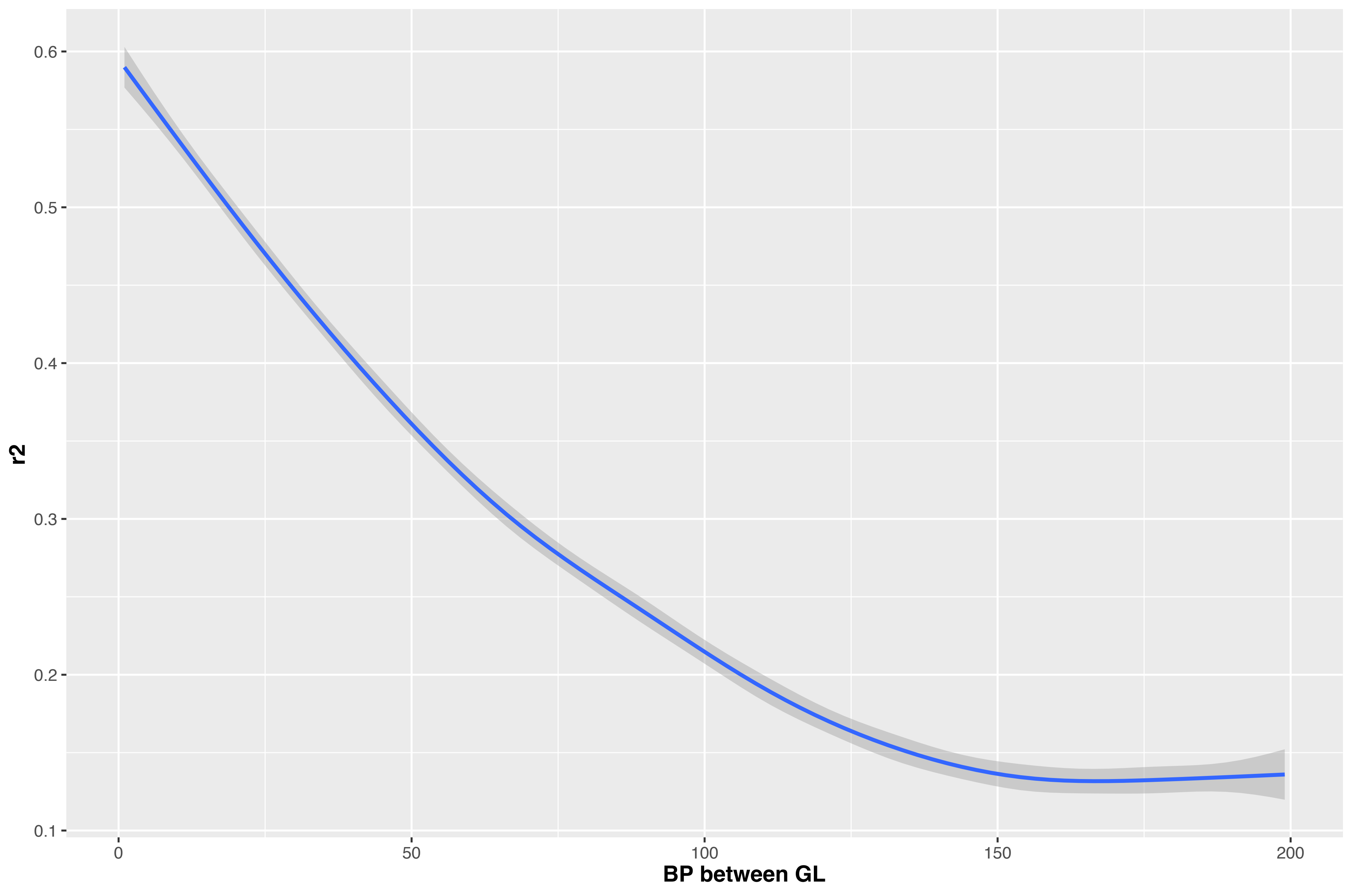

### Figure S3

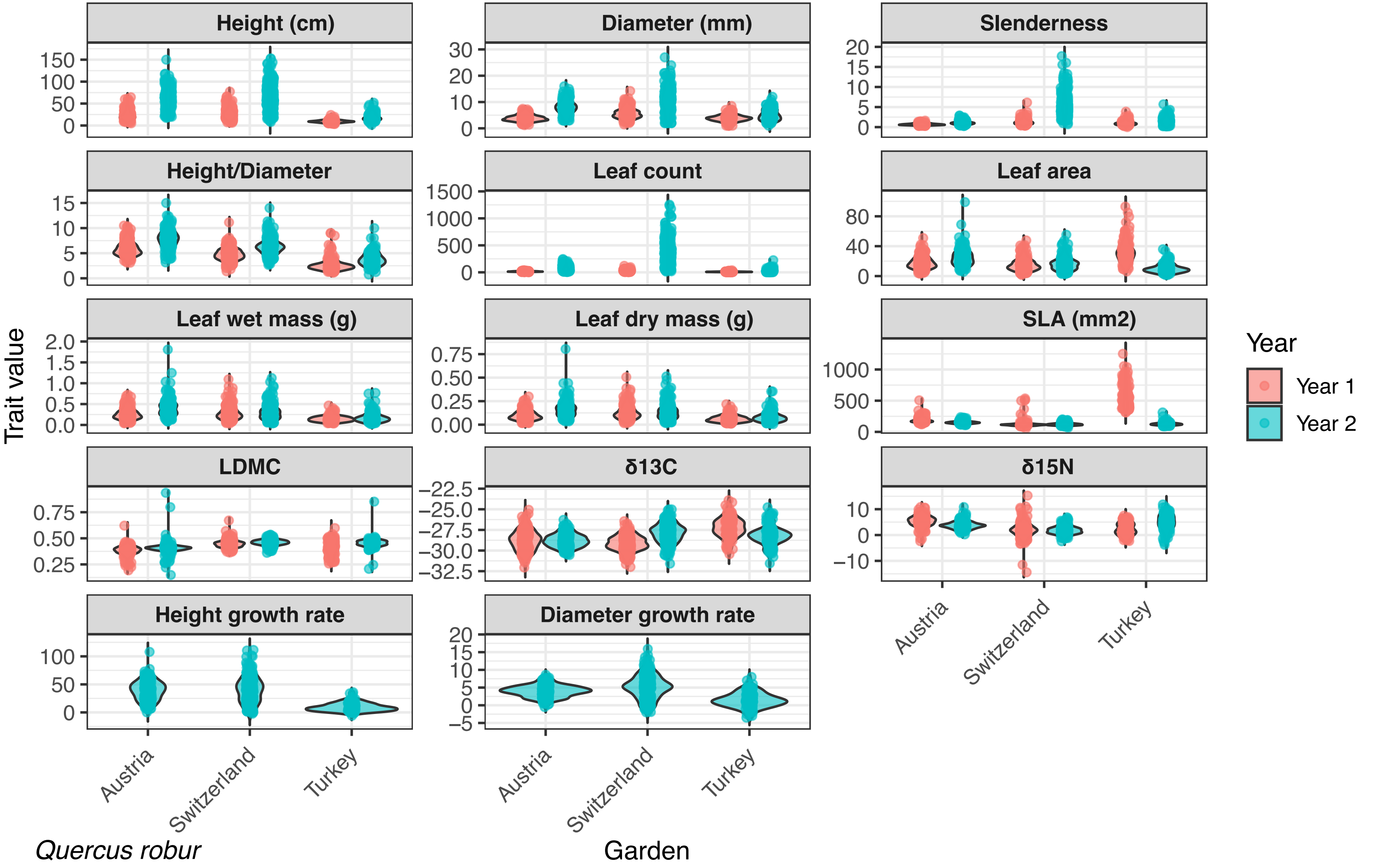

### Figure S4

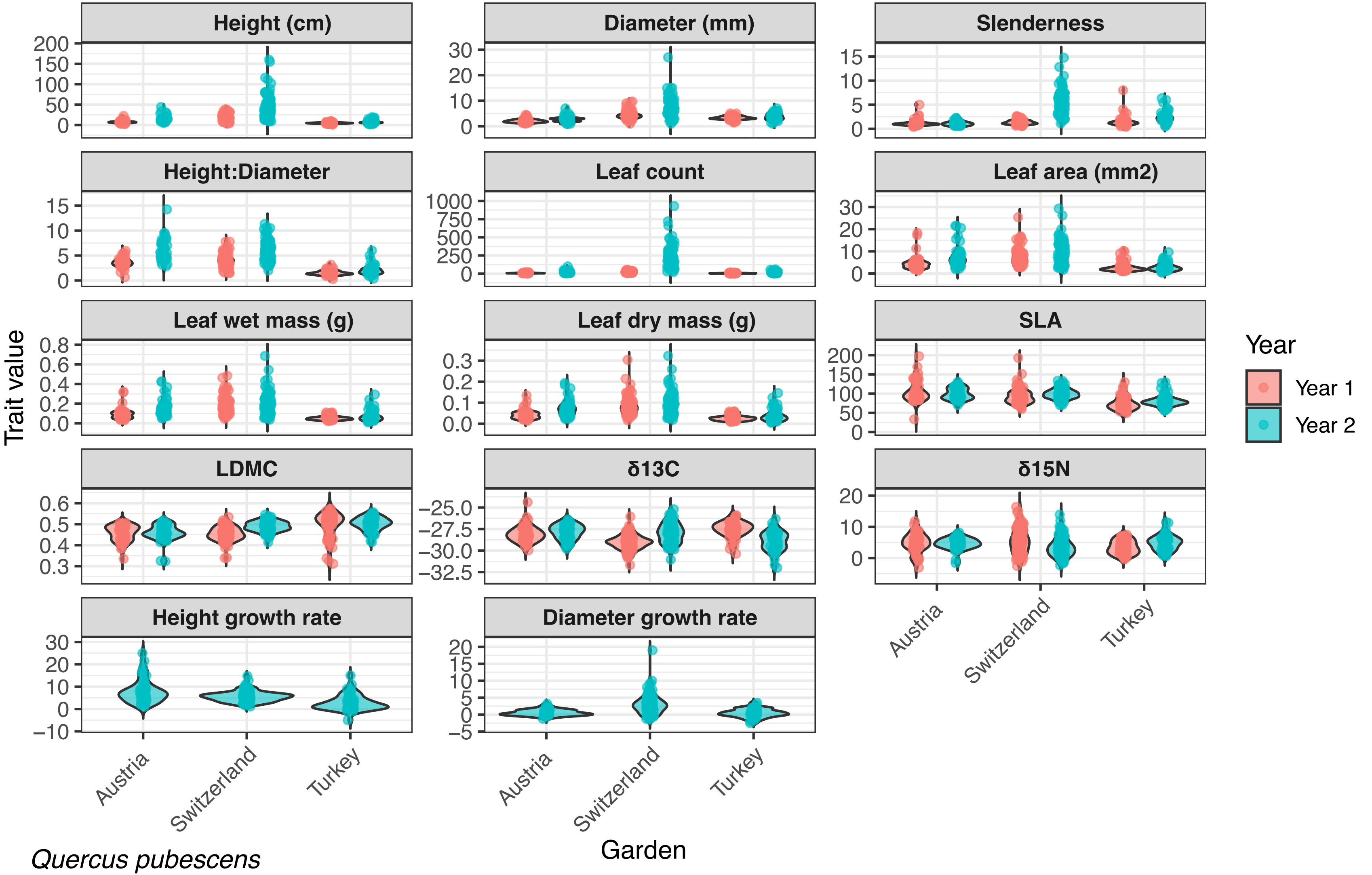

### Figure S5

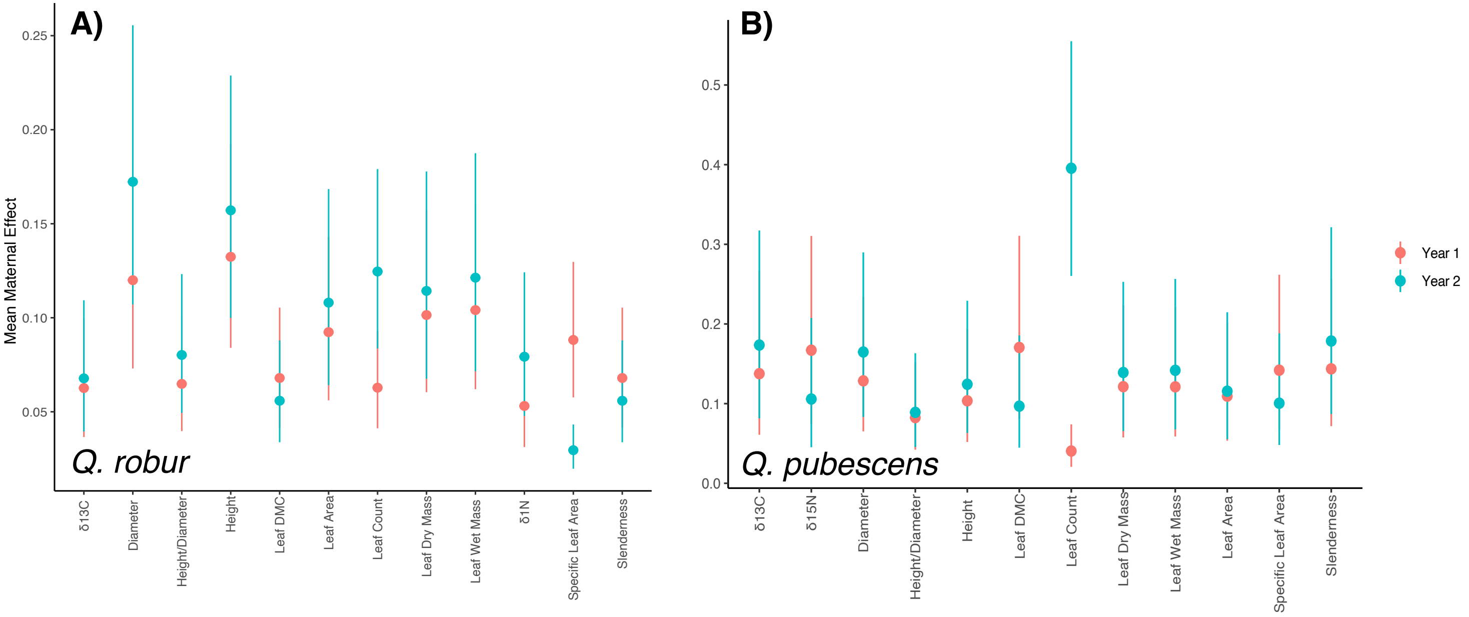

### Figure S6

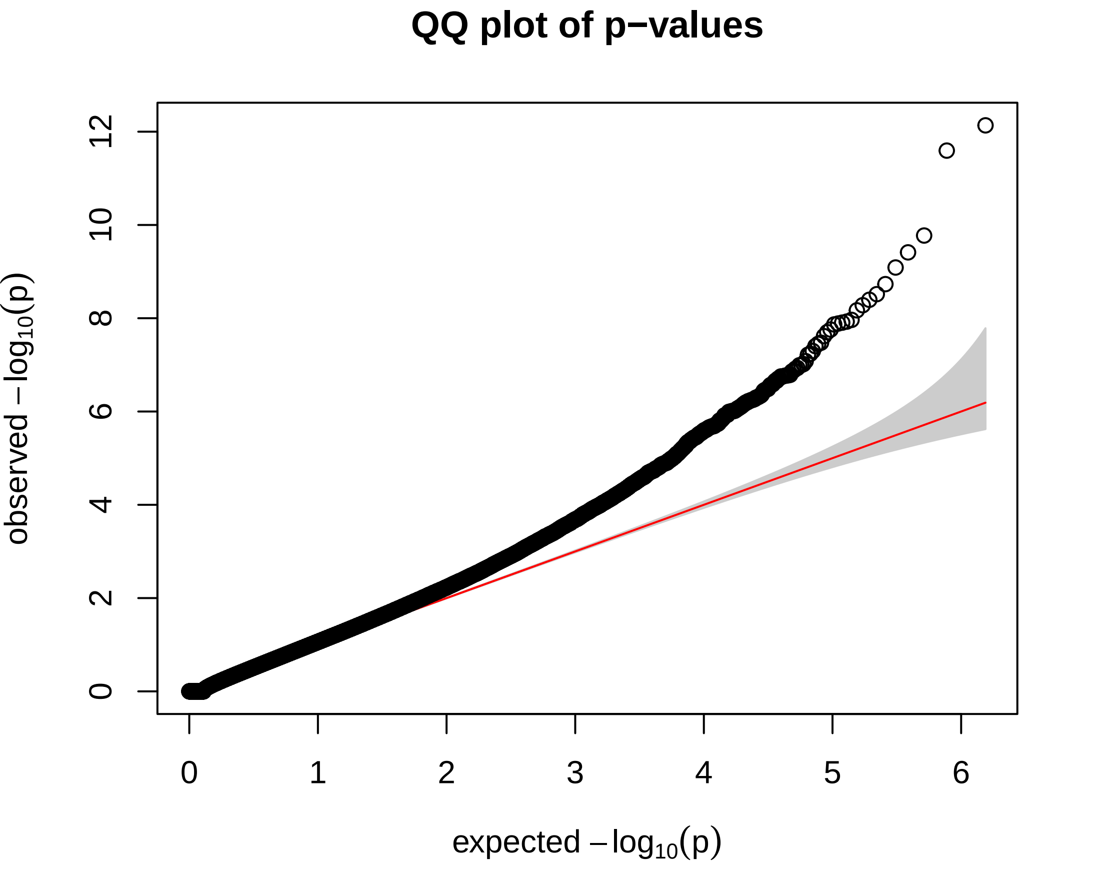

### Figure S7

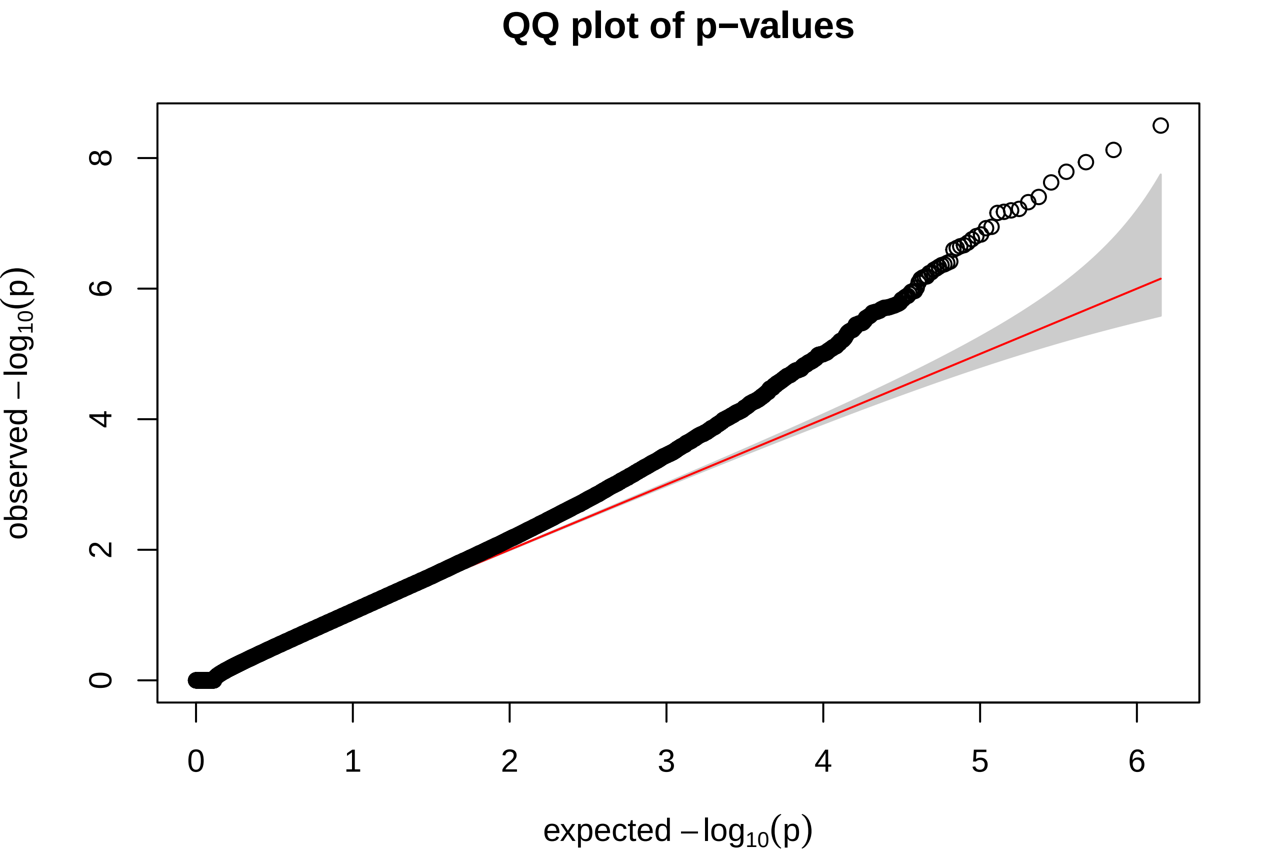

### Figure S8

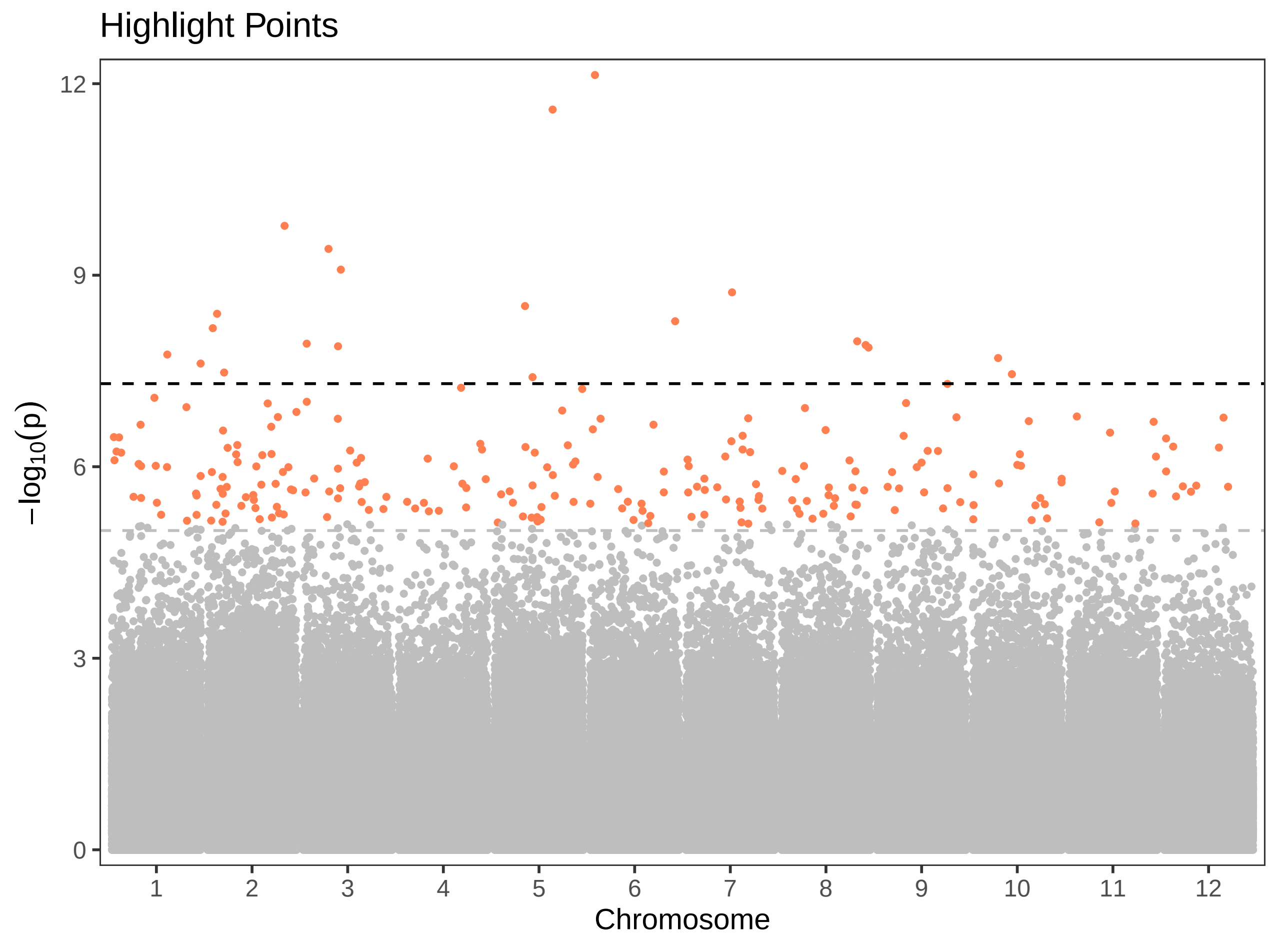

### Figure S9

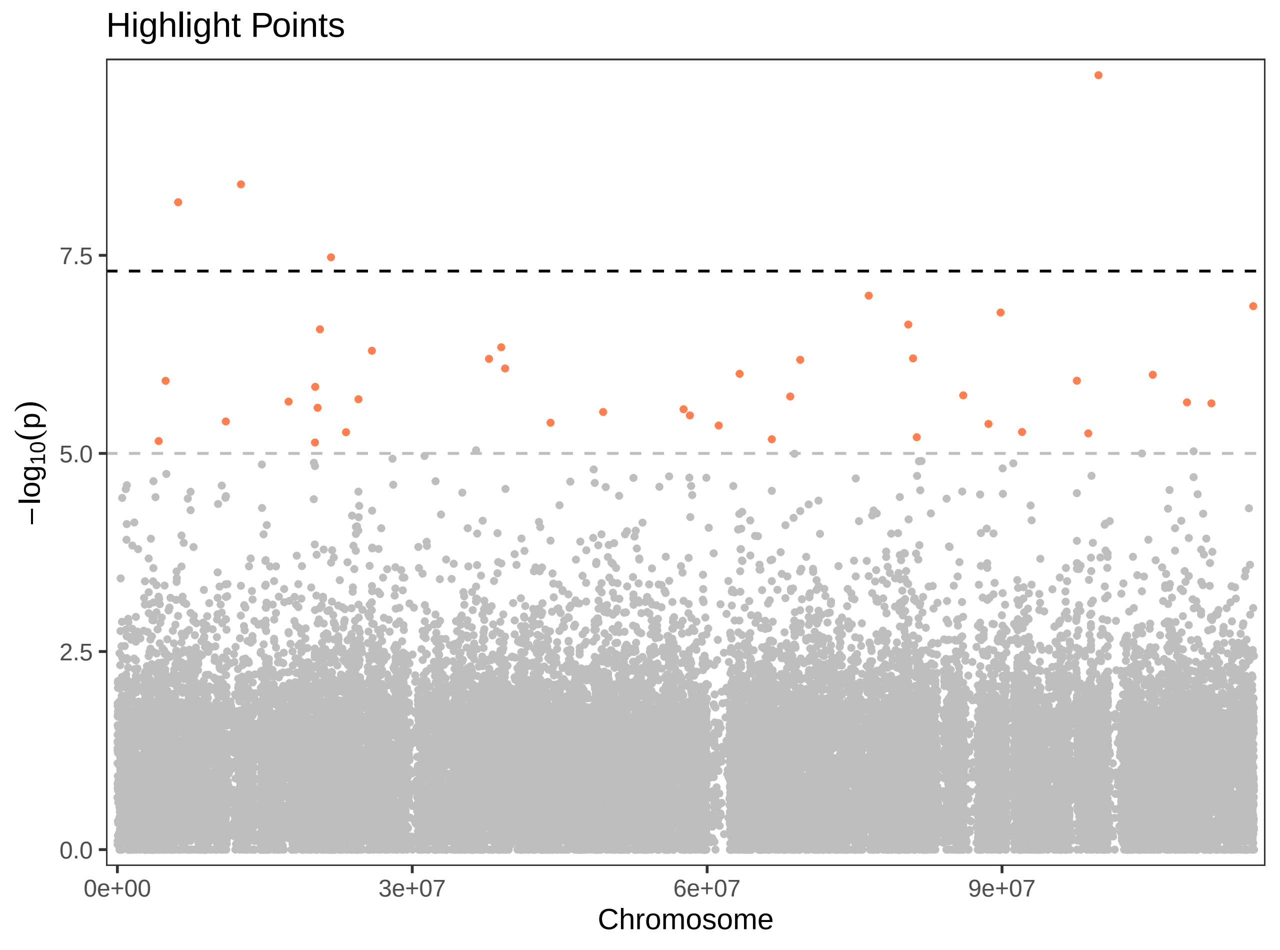

### Figure S10

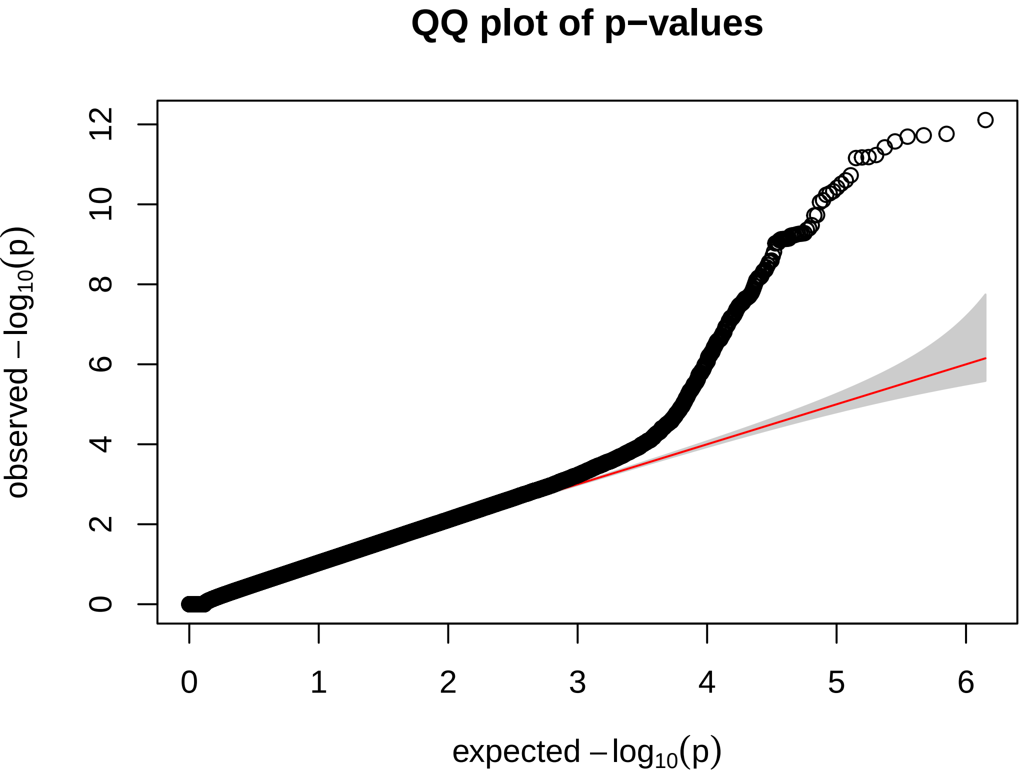

### Figure S11

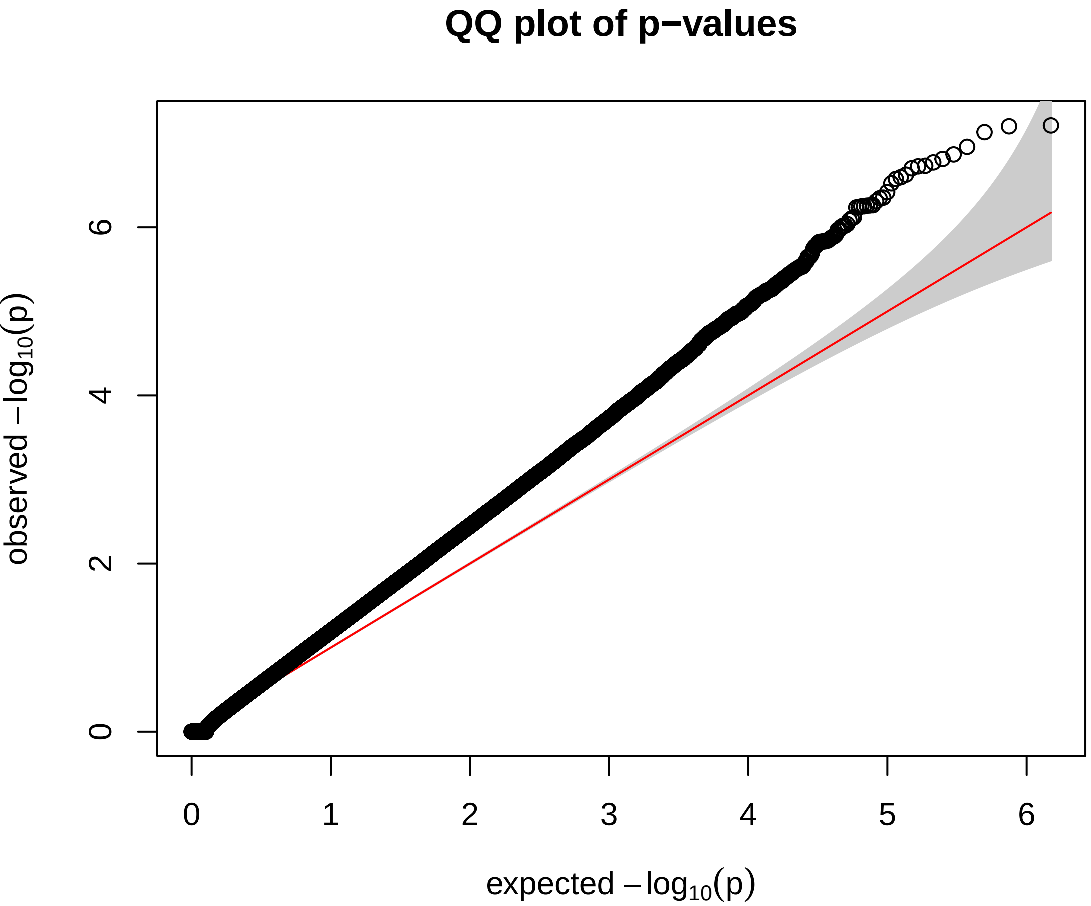
